# Opposing CTCF and GATA4 activities set the pace of chromatin topology remodeling during cardiomyogenesis

**DOI:** 10.1101/2025.10.07.680441

**Authors:** Silvia Becca, Sara Bianchi, Elisa M. Hahn, Kirsten E. Snijders, Lukasz Truszkowski, Anna Krepelova, Francesco Neri, Davide Cacchiarelli, Sasha Mendjan, Elisa Balmas, Alessandro Bertero

**Affiliations:** Molecular Biotechnology Center “Guido Tarone”, Department of Molecular Biotechnology and Health Sciences, University of Torino, Italy; Molecular Biotechnology Center “Guido Tarone”, Department of Life Sciences and Systems Biology, University of Torino, Italy; Telethon Institute of Genetics and Medicine, Pozzuoli, Italy; Institute of Molecular Biotechnology of the Austrian Academy of Sciences (IMBA), Vienna BioCenter, Austria

## Abstract

Reorganization of the three-dimensional chromatin structure is a critical feature of human embryonic development. Yet, the mechanisms regulating integrative remodelling of local structures (e.g., loops) and global architecture (e.g., A/B compartmentalization) remain unclear. Here, we investigate this aspect in the context of cardiomyogenesis, characterized by pronounced B-to-A remodelling of several cardiac-specific genes such as *TTN*. We focus on the roles of the pioneer transcription factor GATA4 and the architectural protein CTCF. Using an inducible knockdown system during human induced pluripotent stem cell differentiation, we show that GATA4 is essential for timely topological activation of key cardiac genes, while partial depletion of CTCF, anticipating physiological downregulation during development, enhances this process. Deletion of a single CTCF binding site on *TTN* leads to modest gene decompaction and transcriptional activation. Bulk and single-cell RNA sequencing of chamber-specific cardiac organoids reveals that loss of GATA4 delays differentiation and sustains proliferation of early cardiomyocytes, whereas premature CTCF depletion accelerates yet alters cardiomyocyte maturation. These findings suggest that CTCF and GATA4 have antagonistic roles on chromatin dynamics during cardiomyogenesis, forming a rheostat that maintains accurate developmental tempo. Disruption of this mechanism may contribute to congenital heart defects caused by mutations in these factors.

## INTRODUCTION

Chromatin architecture dynamically shapes transcriptional programs during development and in human disease^1^, yet the underlying mechanisms remain incompletely understood and are an active area of investigation^2^. During differentiation, pluripotent stem cells (PSCs) undergo extensive chromatin reorganization and transcriptional remodeling to establish lineage-specific gene expression programs. This involves coordinated changes in chromatin accessibility^3^, hi-stone modifications^4^, and three-dimensional (3D) genome organization, a higher-order layer of epigenetic regulation that has recently emerged as a key orchestrator of developmental gene expression^5^. This multi-layered regulation spans from short- and long-range chromatin loops, to topologically associating domains (TADs) hundreds of kilobases long, and up to the seg-regation of megabases-sized chromosomal regions into so-called A (active) and B (inactive) compartments^6^. Together, these structures mediate inter- and intra-chromosomal interactions that shape chromosome territorialization and radial nuclear positioning.

Cardiac development provides a striking example of developmental 3D chromatin remodel-ing, as changes in loops, TADs, A/B compartments, and inter-chromosomal networks facilitate the temporal and spatial control of cardiac gene expression (reviewed in^7^). Disruption of these processes has been linked to genetic dilated cardiomyopathy (DCM) caused by lamin A/C mu-tations^8,9^. Mutations in chromatin regulators are also strongly enriched in *de novo* congenital heart defects (CHD)^10^, underscoring their pathogenic relevance^11^.

Unbiased genome-wide analyses have revealed a remarkable pattern in which large cardiac genes transition from the lamina-associated B compartment to the A compartment within the nu-clear interior, where they engage in higher-order interactions and enhanced transcription^12–14^. These topological changes establish stable gene regulatory networks essential for cardiac lin-eage commitment. However, the molecular mechanisms driving chromatin reorganization in early cardiomyogenesis remain poorly understood, and it is unclear how their dysregulation disrupts cardiac development and function, ultimately leading to CHD and DCM.

Pioneer factors such as GATA4 can access and open closed chromatin regions, enabling recruitment of transcriptional machinery to activate lineage-specific programs^15^. GATA4 is par-ticularly crucial in cardiac development, where it regulates genes controlling mesoderm specifi-cation, first and second heart field formation, and cardiomyocyte maturation^16,17^. Accordingly, mutations in GATA4 have been identified in Mendelian forms of isolated CHD^18,19^.

Architectural proteins such as CTCF (CCCTC-binding factor), a ubiquitously expressed in-sulator, play a key role in defining TADs and mediating enhancer-promoter looping^20^. By part-nering with cohesin, CTCF stabilizes chromatin loops that shape gene regulation of discrete subsets of cell-type specific genes^21,22^. Traditionally viewed as a boundary factor prevent-ing aberrant interactions^23^, CTCF is now recognized as a dynamic regulator of developmental transitions and lineage specification^24,25^. *De novo* CTCF mutations cause a syndromic devel-opmental disorder that includes CHD, implicating its dysfunction in cardiac malformations^26,27^. Moreover, depletion of CTCF in the adult cardiomyocytes leads to heart failure^28,29^

The mechanism of action of GATA4 and CTCF and their combined effects on chromatin reor-ganization in cardiomyogenesis remain largely unexplored. We addressed this question using cardiomyocytes derived from human PSCs and PSC-derived cardiac organoids^30^, which reca-pitulate chamber-specific identities. Our study reveals opposing roles for GATA4 and CTCF, respectively promoting and restraining early cardiac differentiation.

## RESULTS

### The *TTN* locus models B-to-A compartment switching during cardiomyogenesis

We and others previously reported that *TTN*, encoding the large sarcomere protein titin, transitions from the B to A compartment during PSC differentiation into cardiomyocytes (CMs), accompanied by a marked repositioning from the nuclear periphery toward the interior^12,13^. This was evident from shallow-sequenced DNAse Hi-C data from both female RUES2 human embryonic stem cell-derived CMs (hESC-CMs) and male WTC11 human induced pluripotent stem cell-derived CMs (hiPSC-CMs) differentiated for 2 weeks using a biphasic WNT modulation protocol, though with different kinetics (Figure S1A). We observed the same phenomenon by re-analysing deeply sequenced *in situ* Hi-C during a CM differentiation time course of female H9 hESCs up to day 80 (Figure 1A). In all datasets, the compartment switch coincided with progressive reduction of promoter-gene body interactions, indicative of gene unpacking. Quantitatively, interactions between the *TTN* promoter and its gene body decreased by ∼55% from the pluripotent stage to day 80 cardiomyocytes. Thus, *TTN* offers a model to study topological rearrangement that is reproducible across different types of hPSCs, cell lines, and laboratories.

**Figure 1.**
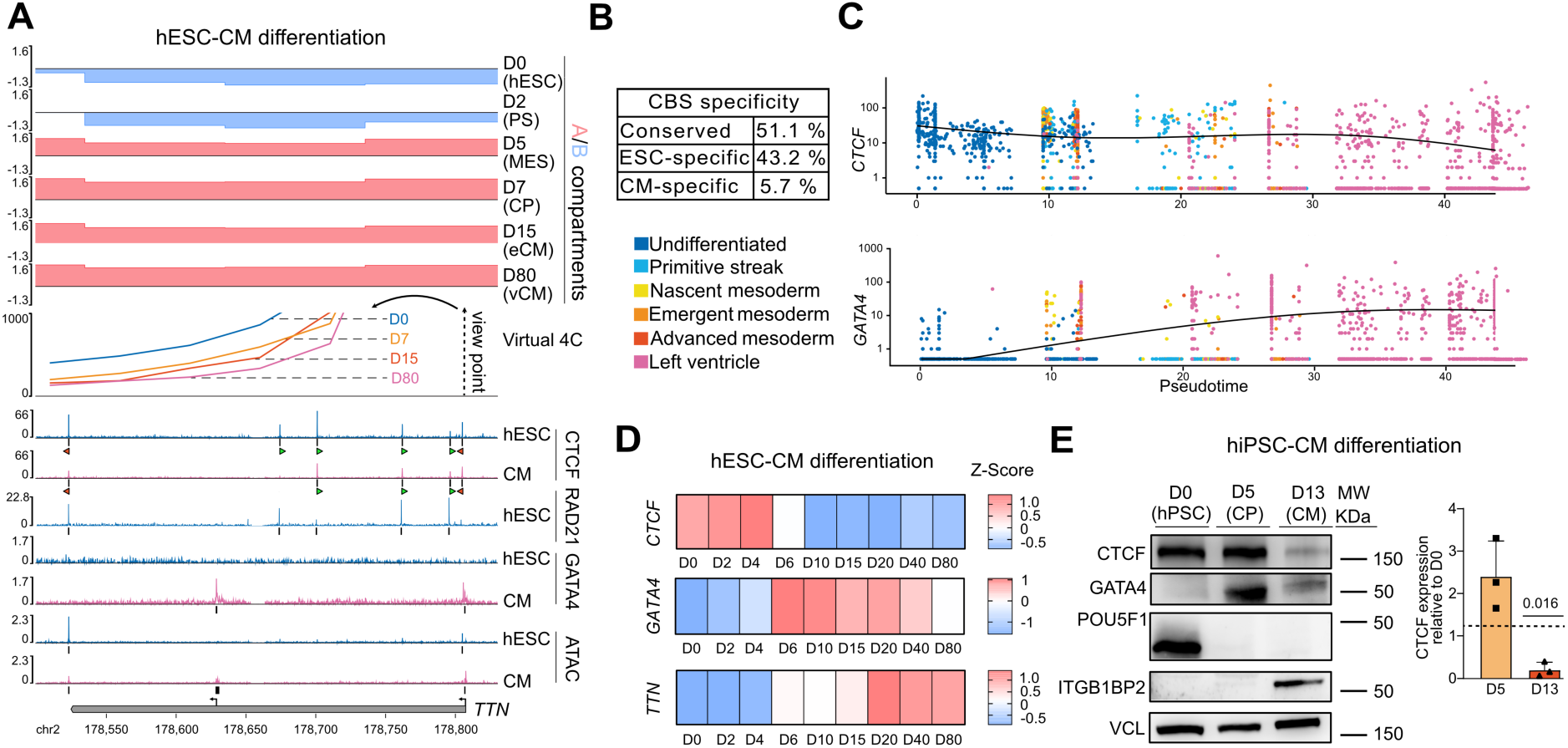
Reciprocal dynamics of the *TTN* locus regulators GATA4 and CTCF. (A) Integrative view of local and global chromatin topology remodelling at *TTN* during H9 hESC-CM differentiation, as assessed by Hi-C (data plotted from quantile normalized average of n = 2 differentiations)^31^, ATAC-seq (data plotted from average of n = 2 differentiations)^13^, and CTCF^32^, RAD21^32^ and GATA4 ChIP-seq^17,33^. Virtual 4C extracted from Hi-C data showing *TTN* DNA interaction with its promoter proximal region. D: day; hESC: human embryonic stem cell; PS: primitive streak; MES: mesoderm; CP: cardiac progenitors; CM: cardiomyocyte; eCM: early CM; vCM: ventricular CM. (B) Percentage of CTCF binding sites (CBS) that are conserved, CM-specific, and hESC-specific^32^. (C) Expression of *GATA4* and *CTCF* from single-cell RNA-seq of human early embryo development^34–36^ (normalized read counts). Cells were developmentally ranked based on pseudotime analysis (Figure S1B). n = 88 pre-implantation embryos, n = 1 gastrulating embryo, and n = 18 fetal heart samples (D) Bulk RNA-seq time course of H9 hESC-CM differentiation^37^. Average z-scores of n = 3 – 4 differentiations. (E) Representative western blot during WTC11 hiPSC-CM differentiation. CTCF expression was normalized to vinculin and to the day 0 sample across n = 3 differentiations; p-value by one-sample t test against *µ* = 1. POU5F1 (OCT4) and ITGB1BP2 (Melusin) are included as established hiPSC and CM markers, respectively^38^.

*TTN* experienced clear changes in chromatin accessibility during CM differentiation: ATAC-seq identified two CM-specific peaks that correlated with ChIP-seq signal for the cardiac pioneer TF GATA4 at the two promoters, one driving full length titin and the other the shorter cronos isoform. In contrast, two regions became less accessible in CMs, correlating with two of the six ChIP-seq peaks for the ubiquitous architectural protein CTCF (Figure 1A). Notably, four out of five putative CTCF binding sites within the *TTN* gene body displayed a convergent orientation with a promoter-proximal CTCF peak. According to the chromatin loop extrusion model, such an arrangement is predicted to favor CTCF-mediated cohesin arrest, stabilizing promoter-gene body interactions that may interfere with transcription initiation and/or elongation^21,39^. Coherently with this hypothesis, RAD21, the chromatin binding subunit of the cohesin complex^40^, co-localizes with five out of the six CTCF ChIP-seq peaks on *TTN* gene body (Figure 1A). Because RAD21 is essential for cohesin loading^41^, this pattern supports the idea that CTCF binding at *TTN* recruits cohesin to form repressive loops that maintain the locus in a compact, transcriptionally inaccessible state. Genome-wide analyses further revealed a substantial reduction in CTCF occupancy during differentiation, with 43.2% of binding sites present in ESCs lost in CMs and only 5.7% newly gained (Figure 1B). This broader decline in CTCF binding suggests that *TTN* exemplifies a more general mechanism by which weakening CTCF–cohesin interactions contributes to chromatin opening during cardiomyogenesis.

Intrigued by these findings, we integrated publicly available single cell RNA sequencing (scRNA-seq) datasets of early human embryo development to determine the expression patterns of *GATA4* and *CTCF* during cardiomyogenesis (Figure 1C, Figure S1B). In line with mouse development^42^, *GATA4* began to be expressed at the nascent mesoderm stage (Figure 1C); this corresponded to ∼5-6 days of directed differentiation towards left ventricle (LV) in monolayer cultures of all three aforementioned hPSC lines (Figure 1D, Figure S1C), and to ∼3.5-4.5 days of WTC11 hiPSC specification into self-organizing LV-, right ventricle-, and atrial-like three-dimensional cardiac organoids (cardioids^30,43^; Figure S1D). Unexpectedly, the increase in *GATA4* expression was mirrored by a decrease in *CTCF* expression, which was approximately halved after mesoderm specification in all models examined (Figures 1B–1C, Figures S1C–S1D). Accordingly, we observed significantly lower CTCF protein levels in WTC11 hiPSC-CMs, the system we selected for further mechanistic studies (Figure 1E).

The decrease in CTCF protein levels may explain its reduced binding on *TTN* and other loci during cardiomyogenesis. Supporting this, the core CTCF binding motifs at *TTN* had lower predicted scores than a constitutively-bound positive control, with the weakest motifs corresponding to the most differentially-bound sites at the 5’ and 3’ end of the gene (Figure S1E). In addition, DNA methylation at the 5’ site was low and remained unchanged throughout differentiation, arguing against methylation as a major factor limiting CTCF binding (Figure S1F). Thus, opposite changes in GATA4 and CTCF expression represent a conserved feature of both *in vivo* and *in vitro* cardiac development and correlate with dynamic occupancy genome-wide.

### GATA4 and CTCF oppositely regulate cardiac B-to-A compartment switching

The correlation between changes in DNA binding and the expression dynamics of GATA4 and CTCF prompted us to test their functional roles in genome topology during cardiomyogenesis. To this end, we generated polyclonal titin-mEGFP reporter hiPSC lines carrying tetracycline (TET)-inducible short hairpin RNAs (shRNAs) for conditional knockdown (KD) of either GATA4 or CTCF, or a scrambled (SCR) shRNA control^44^. We validated at least one shRNA per gene that achieved >70% KD in hiPSC-derived cardiac progenitors (CPs; Figure S2A). Strong de-pletion of GATA4 throughout hiPSC-CM differentiation almost completely blocked cardiac induction, as assessed by loss of cardiac troponin T (cTnT; Figures S2B–S2C). We therefore selected a less effective shRNA to model a more moderate impairment of GATA4, which may better reflect CHD patients carrying heterozygous mutations (Figures 2A–2B). This resulted in a ∼50% decrease in cTnT expression and an ∼8-fold reduction in the proportion of titin-mEGFP^+^ cells, accompanied by a marked drop in the reporter’s median fluorescence intensity(MFI; Figure 2C, Figure S2D). In contrast, CTCF KD during differentiation significantly increased the fraction of titin-mEGFP^+^ cells by more than twofold, with a corresponding rise in MFI, while cTnT levels remained comparable to controls (Figure 2A, Figure S2D).

**Figure 2.**
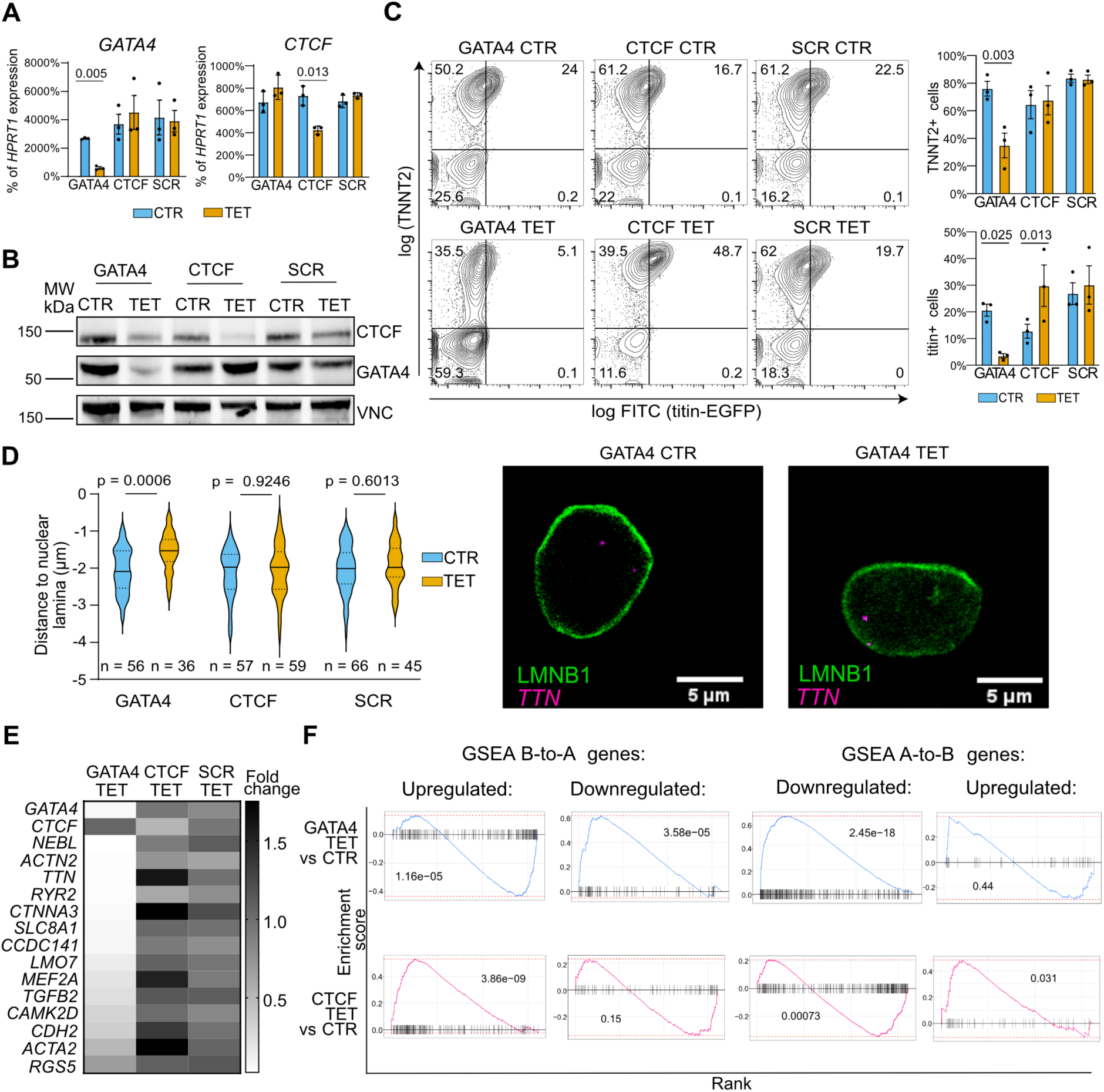
Inducible knockdown of GATA4 or CTCF during hiPSC-CM differentiation. (A) RT-qPCR validation of gene knockdown in day 25 hiPSC-CMs. N = 3 differentiations. p-values by two-way ANOVA with Holm–Sidak corrected pairwise comparisons. CTR: no tetracycline control; TET: tetracycline. (B) Western blot validation of gene knockdown at protein level. VNC: vinculin loading control. (C) Representative flow cytometry quantification of cardiac differentiation efficiency, and quantifications from n =3 differentiations. Statistical analyses as in panel A. (D) ImmunoFISH to quantify the distance of the *TTN* locus from the nuclear lamina (LMNB1). Violin plots report nuclear-normalized distances from the indicated number of cells. p-values one-way ANOVA with with Sidak corrected pairwise comparisons. Representative images correspond to single optical sections from Z-stacks used for 3D volumetric reconstruction. (E) RT-qPCR analyses for a subset of cardiac B to A genes. Average fold-change TET/CTR from n = 3 differentiations. (F) Gene set enrichment analyses (GSEA) of extended cardiac B-to-A and A-to-B gene lists, computed separately for genes up- and downregulated during cardiac differentiation, in bulk RNA-seq from n = 2 differentiations. p-values by Adaptive Monte-Carlo Permutation test.

To connect these expression changes with nuclear topology, we examined the proximity of the *TTN* locus to the nuclear lamina. Following GATA4 KD, *TTN* was more closely associated with lamin B1, consistent with impaired B-to-A compartment transition (Figure 2D). By contrast, CTCF KD did not produce a detectable change in *TTN* positioning (Figure 2D, Figure S2E). Notably, nuclear repositioning of *TTN* was less pronounced in WTC11 hiPSC-CMs than in RUES2 hESC-CMs^13^, consistent with a slower B-to-A transition kinetics for this line (Figure S1A).

To assess whether GATA4 and CTCF have broader roles at other loci undergoing B-to-A transition during cardiomyogenesis, we performed reverse transcription quantitative PCR (RT-qPCR) for 13 such genes we had previously identified during RUES2 hESC-CM differentiation^13^. Most of these genes were downregulated after GATA4 KD, whereas several were upregulated after CTCF KD (Figure 2E). We next broadened the analysis with bulk RNA sequencing (RNA-seq), which was consistent among biological replicates (Figures S2F–S2G). By classifying genomic regions according to their compartment dynamics in H9 ESC–CM differentiation, we found that CM-upregulated genes were significantly enriched in B-to-A regions and depleted in A-to-B regions (chi-squared test). Conversely, CM-downregulated genes showed the opposite pattern(Figure S2H). Gene set enrichment analyses (GSEA) revealed a significant downregulation of CM-upregulated B-to-A genes following GATA4 KD and a significant upregulation of CM-downregulated A-to-B genes. Conversely, CTCF KD led to a significant upregulation of CM-upregulated B-to-A genes and a significant downregulation of CM-downregulated A-to-B genes (Figure 2F). Consistently, expression of genes in the KEGG cardiac muscle contraction pathway was reduced after GATA4 KD but increased after CTCF KD (Figure S2I). Altogether, these results suggest that GATA4 and CTCF function as positive and negative regulators of B-to-A and A-to-B compartment switching.

### CTCF promotes repressive intergenic looping at *TTN*

GATA4 is an established master cardiac TF^17^. By contrast, CTCF has not previously been implicated as a negative modulator of cardiomyogenesis. We therefore focused our attention on this factor, aiming to gain mechanistic insights. Specifically, we perturbed CTCF binding *via* deletion of its cognate DNA sites and examined the effects on chromatin architecture and gene expression (Figure 3A). We first selected the CTCF binding site (CBS) at the gene 5’ end (CBS1) because it is convergent with four other sites within the gene body. As discussed above, the loop extrusion model predicts that CBS1 may stabilize repressive CTCF-mediated loops between the promoter-proximal region and the gene body. We also selected the last site at the gene 3’ end (CBS6), considering that it could contribute to a topologically associating domaing (TAD) boundary isolating *TTN* from the downstream A compartment in hPSCs.

**Figure 3.**
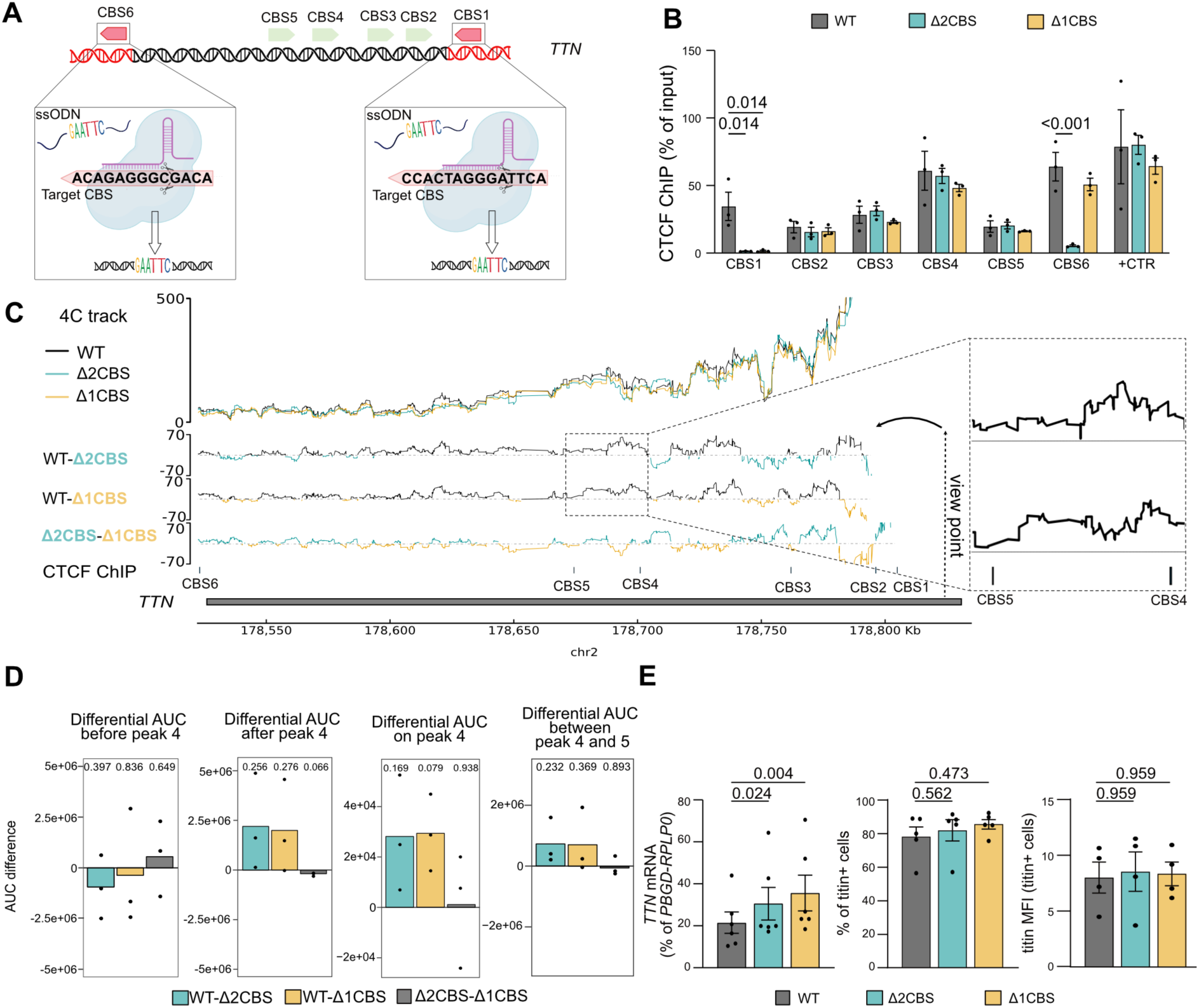
Precise disruption of CTCF binding sites at *TTN*. (A) Schematic of the CRISPR/Cas9 strategy to edit selected CTCF binding sites (CBS) on the *TTN* locus in hiPSCs. The core CTCF binding motif was replaced with an unrelated EcoRI sequence. (B) CTCF ChIP-qPCR in hiPSCs validating genome editing. Enrichment of all *TTN* CBSs was normalized to four negative control regions and compared to a positive control site outside *TTN*. Δ1CBS: homozygous deletion of CBS1; Δ2CBS: homozygous deletion of CBS1 plus compound heterozygous disruption of CBS6; WT: wild-type. n = 3 cultures, p-values by two-way ANOVA with Holm–Sidak corrected multiple comparisons; only significant comparisons are annotated. (C) 4C-seq in hiPSCs using the promoter-proximal region of *TTN* as viewpoint (anchor). The top panel reports raw data, which follows the expected pattern of reduced interactions over genomic distance. To focus on the subtler changes between WT and CBS mutants, the other panels shown the differences of interaction signal between pairs of conditions. Right inset: zoom-in of the CBS4-5 region, demonstrating an increased interaction in the WT versus both mutants. Mean of n = 3 cultures. (D) AUC of the differential 4C-seq signal between pairs of conditions (panel C), for the indicated genomic intervals. p-values by one-sample t-test against *µ* = 0. (E) RT-qPCR of *TTN* mRNA expression (left) and flow cytometry quantification of titin-positive cells and titin median fluorescence intensity (MFI; middle and right) in day 13 hiPSC-CMs. n = 6 differentiations, p-values by two-way RM ANOVA with Holm–Sidak corrected multiple comparisons.

We targeted the core CTCF binding motifs by CRISPR/Cas9-mediated homology-directed repair (HDR) using single-stranded DNA (ssDNA) donor oligos, combined with a co-targeting with selection strategy to increase positive clone recovery^45^ (Figure 3A). We isolated two edited lines: one with homozygous disruption of CBS1 and compound heterozygous editing at CBS6 (Δ2CBS), and another with homozygous editing only at CBS1 (Δ1CBS; Figures S3A–S3B). In the Δ2CBS line, CBS6 carried the intended edit on one allele and a 2-bp deletion on the other (Figure S3C). CTCF ChIP-qPCR in hiPSCs confirmed complete loss of CTCF binding at the targeted sites, including CBS6 in the Δ2CBS line - indicating that the 2-bp deletion sufficed to disrupt CTCF binding, while occupancy at other CBSs remained unaffected (Figure 3B).

We performed circular chromosome conformation capture sequencing (4C-seq) using the promoter proximal region of *TTN* as the viewpoint. The interaction profiles across the *TTN* gene body were very similar for Δ2CBS and Δ1CBS hiPSCs, and each differed in a similar manner from wild-type (WT) controls (Figure 3C). To place these changes in developmental context, we compared our 4C data with virtual 4C generated from published Hi-C of RUES2 hESCs across CM differentiation. Interaction patterns in WT hiPSCs correlated most closely with the pluripotent stage, as expected, whereas Δ2CBS and Δ1CBS clones aligned more closely with cardiac progenitors (Figure S3D). Closer inspection of specific regions revealed two opposing effects in both Δ2CBS and Δ1CBS: promoter interactions increased up to CBS2, but decreased beyond CBS4, particularly between CBS4 and CBS5 (Figures 3C–3D, Figures S3E–S3F). While these differences did not reach significance, the trends suggest that CBS1 deletion is sufficient to mimic some developmental chromatin topology changes by disrupting long-range intergenic looping between the *TTN* promoter and gene body, favoring shorter-range interactions typical of a less constrained chromatin polymer.

We next asked whether these architectural changes influenced *TTN* expression. RT-qPCR revealed a significant increase in *TTN* mRNA by 50-75% in both Δ2CBS and Δ1CBS hiPSC-CMs, despite comparable efficiency of CM specification (Figure 3E). Titin protein levels, however, remained unchanged, likely due to compensatory mechanisms buffering the modest gene expression increase (Figure 3E). The absence of further changes in Δ2CBS compared to Δ1CBS is consistent with the 4C data and indicates that CBS6 is redundant in this context.

Overall, deletion of a single CTCF binding motif produced only moderate effects on *TTN* chromatin organization and transcription, consistent with the redundancy of CTCF sites at large loci^46,47^. Nevertheless, these findings support a model in which CTCF-mediated intergenic looping at *TTN* contributes to transcriptional restraint, providing a mechanistic link to the broader derepression of B-to-A genes observed upon partial depletion of CTCF.

### GATA4 and CTCF exert antagonistic control over second heart field development

To explore the physiological relevance of GATA4- and CTCF-dependent chromatin topology, we turned to a more complex three dimensional (3D) model: self-assembling, multi-lineage cardioids^30,43^. Using scRNA-seq, we validated protocols to generate WTC11 hiPSC-derived 3D cardioids representing the three primary heart chambers: the first heart field (FHF)-derived left ventricle (LV), and the second heart field (SHF)-derived right ventricle (RV) and atria (A; (Figure S4A).

Each cardioid type contained the major cardiac cell populations, including cardiomyocytes (*TNNT2*^+^ and *TTN*^+^), fibroblasts (*COL3A1*^+^), and endothelial cells (*ERG*^+^), while maintaining chamber-specific identities (e.g., *IRX4*^+^ LV CMs, *IRX1*^+^ RV CMs, and *N2RF2*^+^ A CMs; Figures S4A–S4B). In addition, we identified proliferating cells (*MKI67*^+^), mesodermal-stage cells (*MESP1*^+^), and small fractions of undifferentiated (*POU5F1*^+^) and epithelial-like (*KRT19*^+^) cells (Figure S4A).

We investigated the effects of GATA4 and CTCF silencing in this model by differentiating inducible KD hiPSCs into FHF- and SHF-derived cardioids for 7.5 days. RT-qPCR indicated that KD did not alter chamber identity (Figure S5A). GATA4 KD cardioids showed upregulation of both FHF and SHF cardiac progenitor markers (*HAND1*, *HAND2*, *NKX2-5*), together with increased expression of the fibroblast marker *COL1A1* (Figure S5A). In contrast, CTCF KD caused downregulation of *COL1A1* (Figure S5A). Consistent with findings in monolayer hiPSC-CMs, *TTN* expression was decreased by GATA4 KD in all cardioid types and increased by CTCF KD in RV and A cardioids, though intriguingly not in LV ones (Figure 4A). The fraction of titin^+^ cells largely mirrored the mRNA changes, though differences were more modest (Figure 4B), possibly because titin expression was more accelerated and homogeneous in cardioids compared with monolayer hiPSC-CMs (Figure 2C).

**Figure 4.**
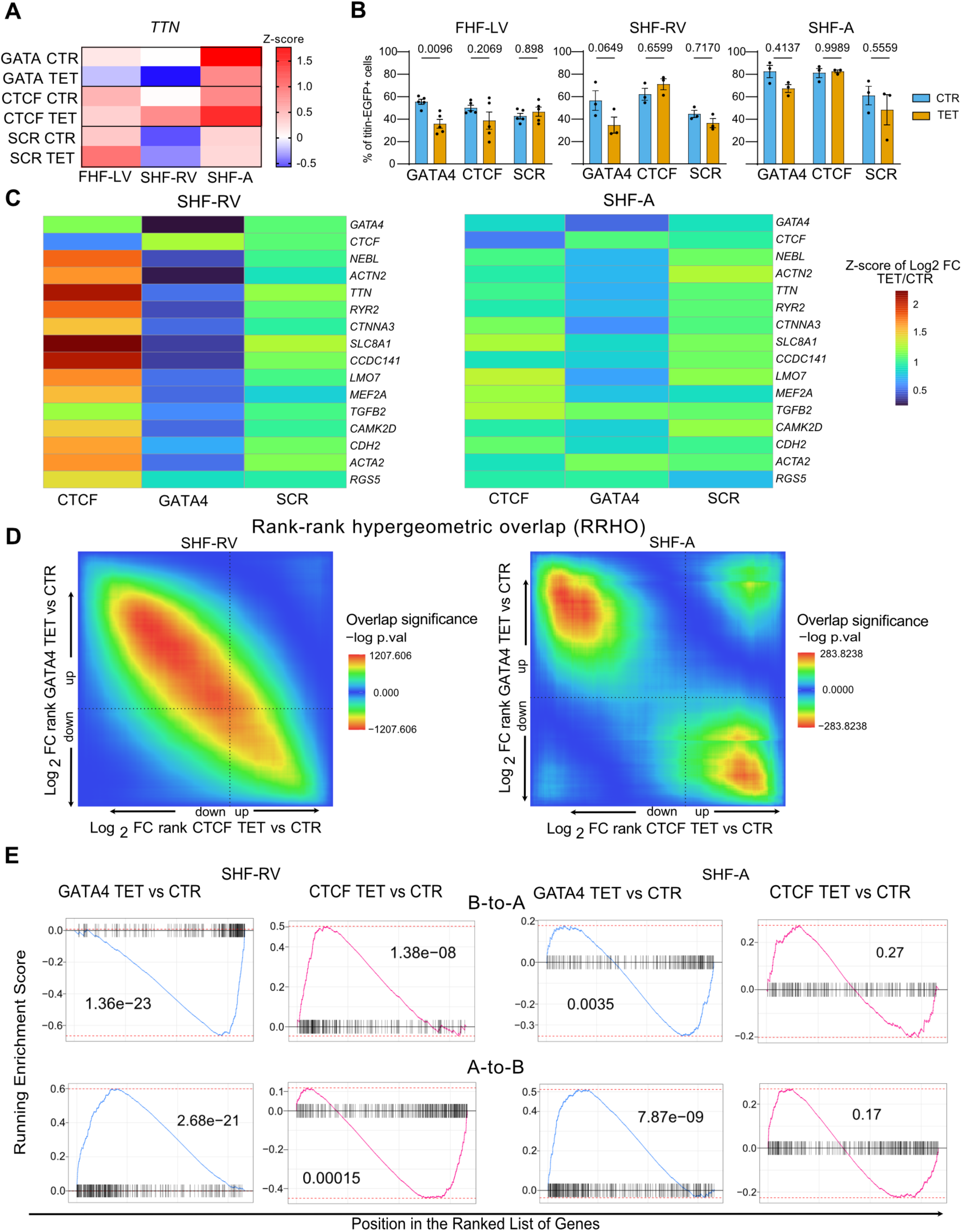
Inducible knockdown during FHF and SHF cardioid differentiation. (A) RT-qPCR of day 7.5 inducible knockdown cardioid. n = 3 differentiations. Data are shown as average z-scores. FHF-LV: first heart field-derived left ventricle; SHF-RV: second heart field-derived right ventricle; SHF-A: second heart field-derived atria. (B) Flow cytometry analyses of cardioids described in panel A. p–values by two-way RM ANOVA with Holm-Sidak corrected pairwise comparisons. (C) Bulk RNA-seq in SHF-RV and SHF-A cardioids. Expression of selected B-to-A cardiac genes (refer to Figure 2E), shown as z-scored log_2_ fold change (TET/CTR). n = 2 differentiations. (D) Rank-rank hypergeometric overlap (RRHO) comparing RNA-seq results for GATA4 KD *versus* CTCF KD. p-values by hypergeometric tests across thresholds of differential gene expression ranks. (E) GSEA for CM-upregulated B-to-A and CM-downregulated A-to-B genes. p-values by Adaptive Monte-Carlo Permutation test.

Because the effects of GATA4 and CTCF silencing were more evident in SHF derivatives, we performed bulk RNA-seq on day 7.5 RV and A cardioids in biological duplicate, which showed good reproducibility and clustering by KD type (Figure S5B). Expression analysis of B-to-A genes revealed a general downregulation after GATA4 KD and an upregulation after CTCF KD in RV organoids, while effects in atrial organoids were modest (Figure 4C). We next compared global transcriptional changes between conditions using rank-rank hypergeometric overlap (RRHO) analysis^48^, which demonstrated a strong anti-correlation between GATA4 and CTCF KD profiles, particularly in RV organoids (Figure 4D). To further probe these transcriptional profiles, we performed GSEA comparing KD vs control in both chamber types. In RV cardioids, GATA4 KD produced a pronounced downregulation of gene sets related to sarcomere organization, cardiac muscle cell development, and heart process and contraction, alongside an upregulation of cell cycle and DNA replication terms. Conversely, RV CTCF KD displayed significant enriched for upregulated genes associated with muscle contraction and development (Figure S5C). Finally, GSEA of CM-upregulated B-to-A and CM-downregulated A-to-B genes showed significant delay of maturation after GATA4 KD in both RV and A cardioids, with a significant downregulation of B-to-A genes expression and upregulation of A-to-B genes expression (Figure 4E). Conversely, CTCF KD accelerated transcriptional maturation, significantly upregulating B-to-A genes and downregulating A-to-B genes in RV cardioids (Figure 4E). Together, these observations indicate that GATA4 and CTCF exert opposing control over B-to-A and A-to-B gene expression and, more broadly, cardiomyocyte development of the SHF, particularly the RV.

### GATA4 and CTCF regulate the timing and quality of first heart field differentiation

Because CTCF silencing produced limited changes in LV cardioids when assessed in bulk, we investigated this chamber type with higher resolution using scRNA-seq. To capture potential early differences, we analyzed LV cardioids at both day 4.5 and day 7.5 of differentiation, each in biological duplicate. We reasoned that LV differentiation may progress so rapidly that early transcriptional changes could be compensated by later stages. This may explain some of the differences between LV *versus* and SHF-derived cardioids, which in our protocol enter the final stage of differentiation one day later, to mimic the slower timing of SHF development^30,43^.

Day 4.5 LV cardioids from inducible KD lines maintained in control conditions were highly similar to each other and to TET-treated SCR controls, as expected (Figure S6A). These cardioids were composed primarily of CMs (∼50%) and their progenitors (∼10%), fibroblasts (∼25%), and a smaller fraction of earlier developmental populations (Figures 5A–5B and Figure S6B). Biological replicates also showed strong reproducibility (Figure S6A). GATA4 KD resulted in an expansion of undifferentiated and mesodermal cells accompanied by a reduction in CMs (Figure 5B). In contrast, CTCF did not alter the overall CM fraction. Notably, both perturbations markedly altered CM identity, generating clusters that were virtually unique to GATA4 or CTCF KD. Pseudotime analysis ranked these CM populations as the least mature (GATA4 KD) and most mature (CTCF KD), respectively, with the trajectory correlating with increasing expression of B-to-A genes such as *TTN* and *RYR2* (Figure 5C). Milo-based differential abundance testing^49^ confirmed that the shifts in cell populations after GATA4 and CTCF KD were robust across replicates and at the neighborhood level (Figure 5D).

**Figure 5.**
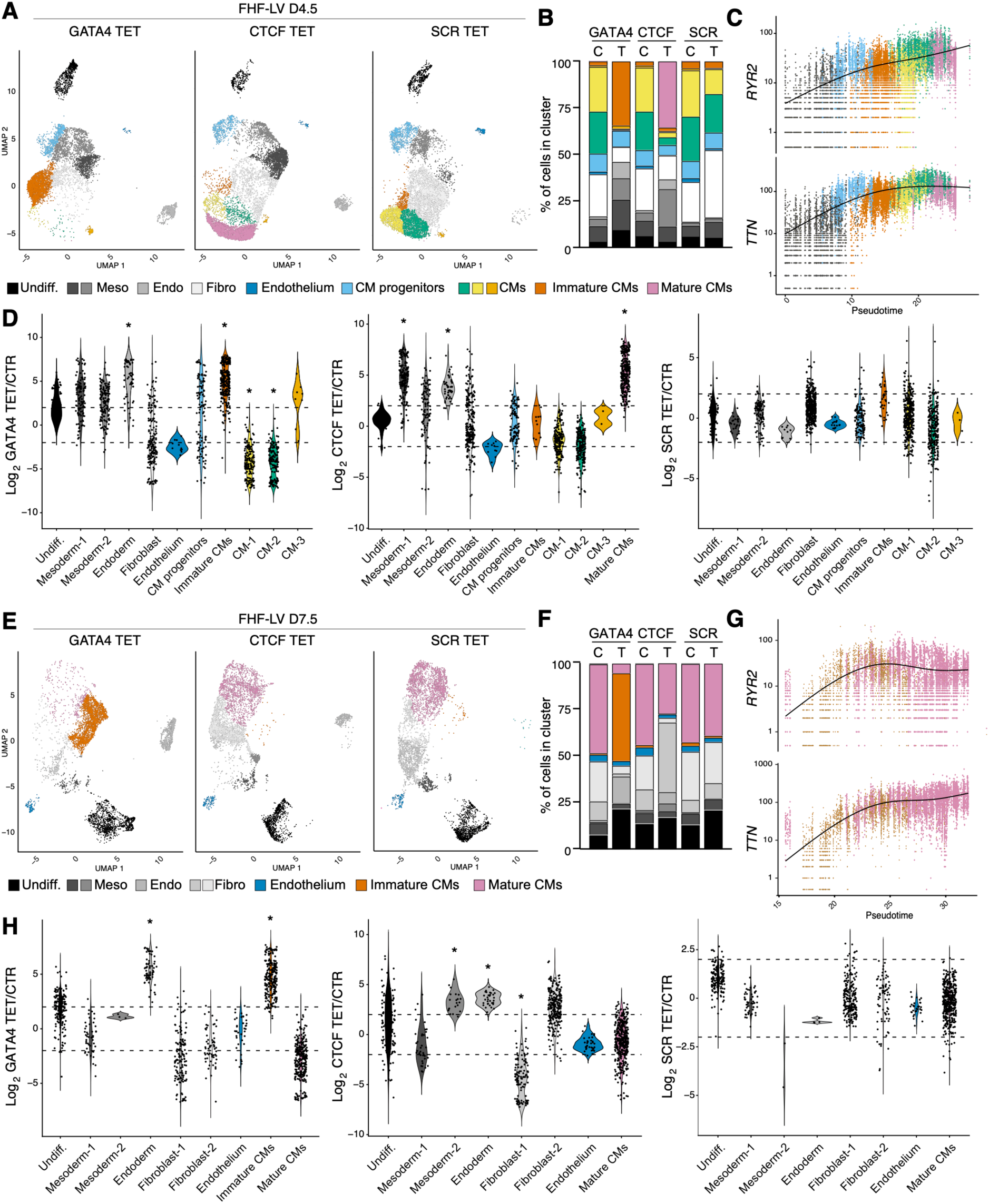
Single-cell RNA-seq of differentiating inducible knockdown LV cardioids. (A) UMAP representation of scRNA-seq data from two differentiations of day 4.5 FHF-derived LV cardioids with inducible knockown. (B) Cell type composition, shown as percentage of cells per cluster. C: control; T: tetracycline. (C) Gene expression in cardiac progenitors and CM subtypes at day 4.5, ranked by pseudotime. (D) Differential cluster abundance analysis at day 4.5 using graph neighborhoods (KNN). Significantly different clusters (*) are identified by an absolute median logFC > 0.05, a fraction of significant cells > 0.75, and a minimal spatial FDR < 0.05. (E) UMAP representation of scRNA-seq data from two differentiations of day 7.5 FHF-derived LV cardioids with inducible knockdown. (F) Cell type composition, shown as percentage of cells per cluster. (G) Gene expression in cardiac progenitors and CM subtypes at day 7.5, ranked by pseudotime. (H) Differential cluster abundance analysis at day 7.5 using graph neighborhoods (KNN). Significant changes defined as described in panel D..

By day 7.5, control LV cardioids again showed strong similarity (Figure S6A). Their overall cellular composition was comparable to day 4.5, though CMs appeared more mature, as ex-pected (Figures S6C–S6D). At this stage, GATA4 KD produced a distinct cluster of immature CMs with reduced B-to-A gene expression (Figures 5E–5G). In contrast, CTCF KD no longer yielded a distinct CM cluster but instead showed a modestly lower CM fraction, consistent with flow cytometry results (Figure 4B). This reduction was accompanied by the expansion of a poorly defined population co-expressing fibroblast and cardiac markers (Figure 5H and Fig-ure S6C). Together, these findings indicate that while CTCF functions as a general brake on CM differentiation, it is nonetheless required for proper LV chamber development.

### GATA4 promotes cardiomyocyte cell cycle exit

To assess the functional consequences of GATA4 and CTCF silencing on cardioid morphogene-sis, we quantified LV cardioid size from day 2.5 to 7.5 of differentiation. GATA4 KD cardioids be-came significantly larger, whereas CTCF KD had no marked effect (Figure 6A and Figure S7A). Because scRNA-seq had indicated an accumulation of immature cardiomyocytes in GATA4 KD cardioids, we hypothesized that this enlargement might reflect increased proliferation. To test this, we simultaneously measured DNA synthesis and DNA content in day 4.5 LV cardioids, an assay commonly used to assess cell cycle distribution. GATA4 KD resulted in a significantly lower proportion of titin^+^ CMs in G1 and a higher fraction in S and G2 phases (Figures 6B–6C and Figure S7B). Non-CMs also appeared slightly more proliferative (Figure S7C). Histological staining of MKI67, a marker of DNA synthesis, supported these findings (Figure S7D).

**Figure 6.**
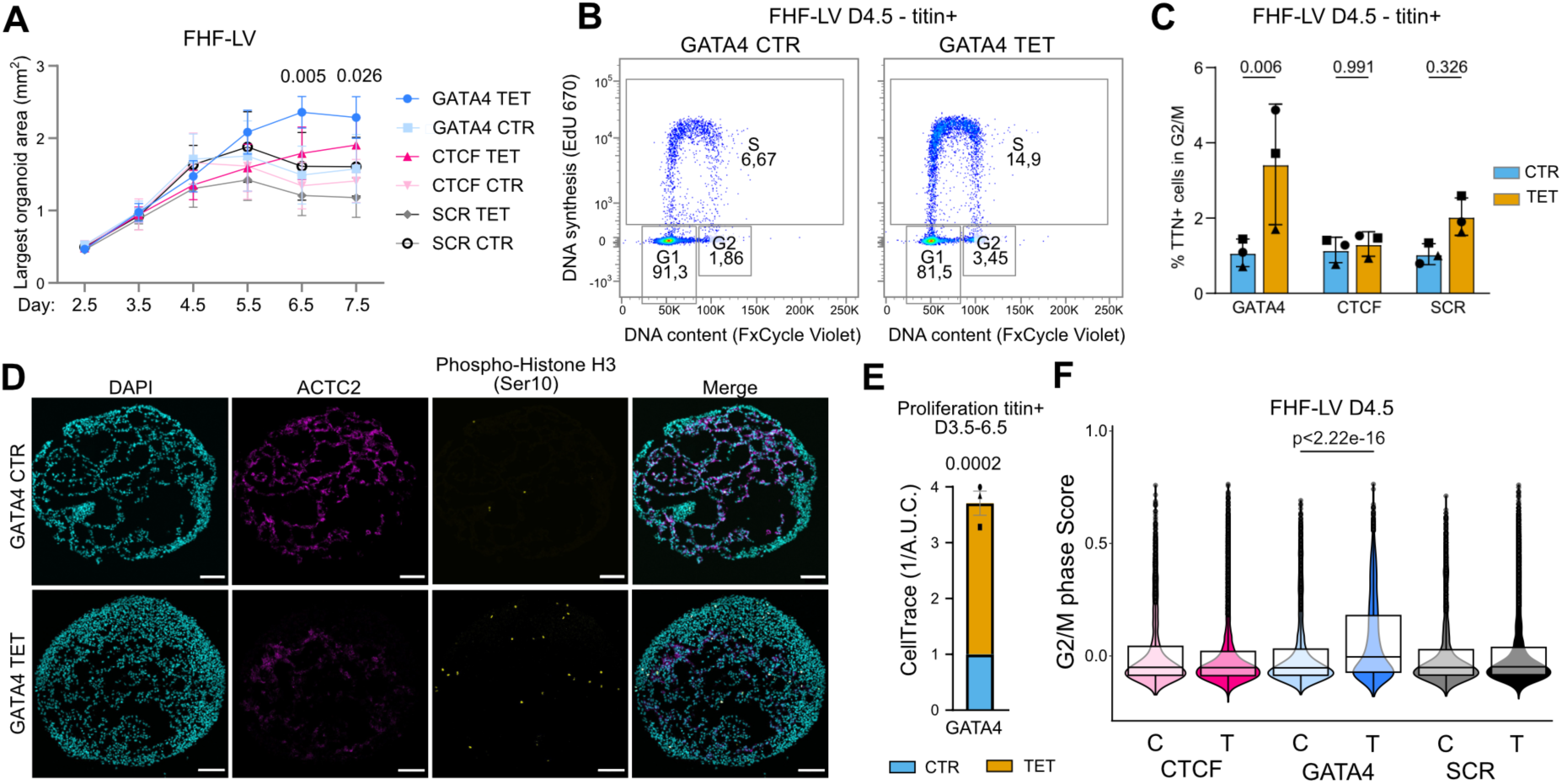
Proliferation dynamics in inducible knockdown LV cardioids. (A) Quantification of LV cardioid size progression. n = 3 differentiation, 8 cardioids each. p-values by two-way ANOVA with Holm-Sidak corrected pairwise multiple comparisons. (B) Representative flow cytometry plots of cell cycle distribution in titin^+^ CMs at day 4.5, based on 5-ethynyl-2′-deoxyuridine (EdU) incorporation and DNA content. (C) Quantification of the experiment shown in panel B. n = 3 differentiations, p-values by two-way RM ANOVA with Holm-Sidak corrected pairwise multiple comparisons. (D) Immunofluorescence of day 4.5 LV cardioids stained for *α*-actinin 2 (ACTC2), nuclei (DAPI) and phospho-histone H3. Scale bar: 100 µm. (E) Proliferation assay with CellTrace dye. Cardioids were labelled at day 3.5 and analyzed by flow cytometry at day 6.5. Data show fold dilution of CellTrace signal in titin^+^ CMs, normalized to baseline (day 3.5). n = 3 differentiations, p-value by one-sample t-test against *µ* = 1. (F) Bioinformatic inference of G2/M phase from scRNA-seq data of day 4.5 cardioids. p-value by Wilcoxon test.

Because maturing CMs can undergo endoreplication leading to polyploidy, DNA synthesis and G2-like DNA content alone cannot conclusively demonstrate proliferation. We therefore assessed phospho-histone H3, a more specific marker of late G2/M, which was also increased in GATA4 KD cardioids (Figure 6D). To obtain a direct measure of cytokinesis, we labeled car-dioids at day 3.5 with a cell tracer and monitored its dilution through day 6.5. This confirmed an increased proliferation rate, particularly in titin^+^ CMs (Figure 6E and Figures S7E–S7F). Finally, bioinformatic inference of cell cycle phase from scRNA-seq data further supported increased proliferation of GATA4 KD CMs (Figure 6F). Altogether, these data show that GATA4 promotes cardiomyocyte cell cycle exit, whereas CTCF silencing, despite promoting cardiac maturation, has little effect on proliferation, underscoring their distinct roles in LV development.

## DISCUSSION

It is well established that genome architecture is highly dynamic during embryonic development. Here, we leveraged monolayer cardiomyocyte differentiation and chamber-specific multilineage 3D organoids to shed light on the mechanisms driving lineage-specific gene activation during a clinically relevant developmental trajectory. Indeed, CHD is the most common birth defect, affecting ∼1% of all live births and ∼0.4% of adults^25^. Our working model is that GATA4 plays a pivotal role in the early chromatin remodeling required to activate cardiomyogenesis, while CTCF puts a brake on progression towards cardiomyocyte maturation. We propose that these concerted activities ensure a sufficient pool of early cardiomyocytes before cell cycle exit and maturation, and that disruption of either checkpoint compromises chamber formation (Figure 7).

**Figure 7.**
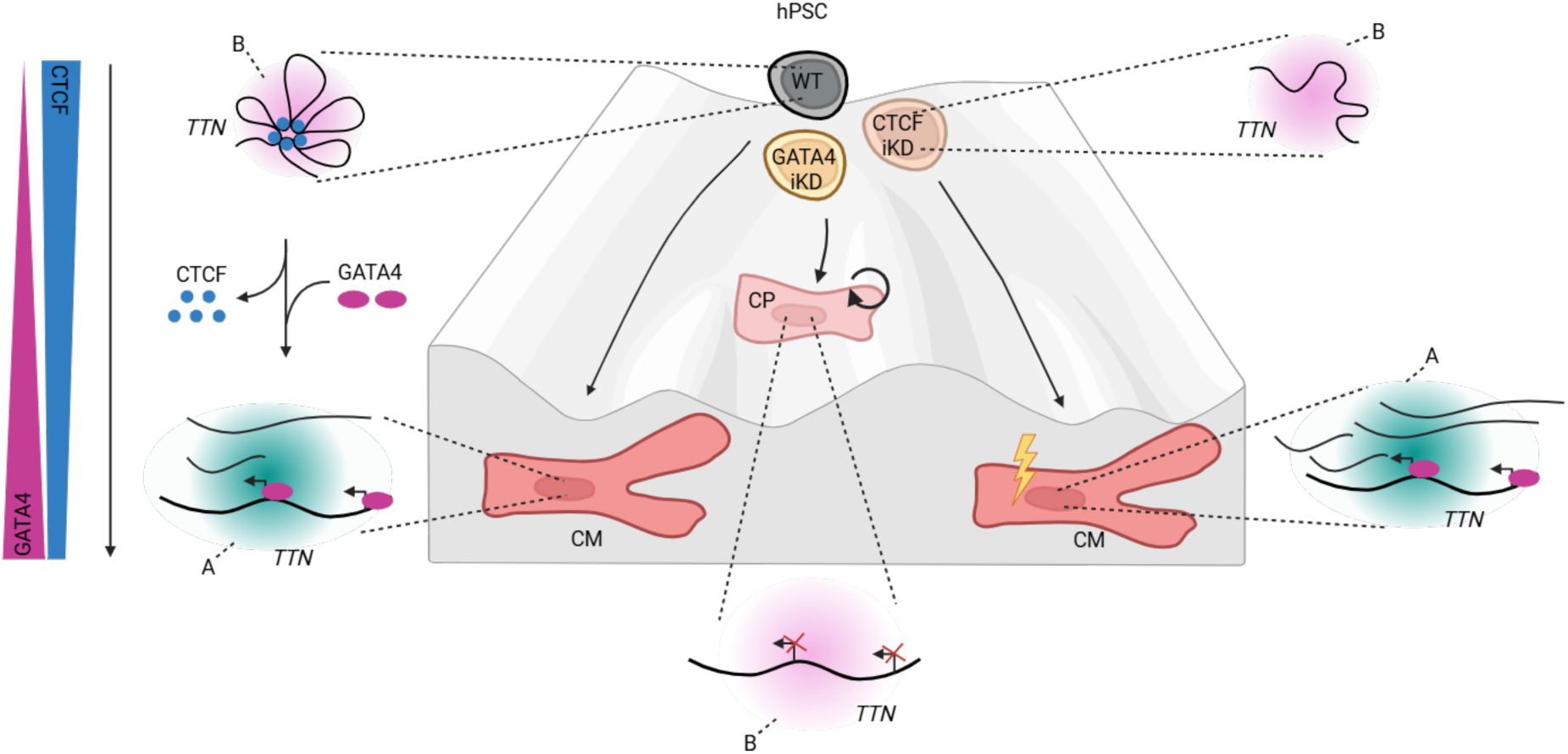
Working model of GATA4 and CTCF function in cardiomyocyte differentiation. During cardiomyocyte differentiation, GATA4 acts as a pioneer transcription factor that drives decompaction and transcriptional activation of cardiac genes undergoing B-to-A switching, exemplified by *TTN*. Reduced GATA4 expression leaves cardiac progenitors in an immature, proliferative state, marked by low cardiac gene expression and persistent association of these loci with the nuclear lamina. Conversely, CTCF, which is highly expressed in pluripotent cells, restricts cardiac gene accessibility and prevents premature activation. As differentiation progresses, declining CTCF levels allow timely chromatin remodeling. Premature CTCF depletion facilitates cardiac maturation but disrupts proper chamber morphogenesis.

In line with a previous hypothesis on *ENPP3* regulation in HEK239T cells^50^, our data indicate that CTCF contributes to the compact organization of *TTN* in the undifferentiated state through repressive loops formed at convergent binding sites in its 5’ region. These loops restrict *TTN* accessibility when not required and may represent a general mechanism for other B-to-A cardiac genes regulated by CTCF. During differentiation, CTCF levels decrease to about half of those in hPSCs, which may be sufficient to release weaker binding sites at *TTN*. Other “epigenetic brakes” have also been described in differentiating cortical neurons^51^, suggesting a broader role for such mechanisms in development.

Reduction in CTCF binding coincides with activation of the pioneer factor GATA4, rendering *TTN* accessible to transcription at the appropriate time. We previously observed that GATA4 binding is enriched at newly accessible chromatin regions in both cardiac progenitors and cardiomyocytes, whereas NKX2-5 and TBX5, key cardiac TFs which lack pioneer activity, are enriched only at cardiomyocyte-specific accessible sites^13^. In TBX5 haploinsufficient cardiomyocytes, reduced TBX5 dosage increased CTCF binding at cardiac genes^14^. Together, these findings support a model in which GATA4-triggered chromatin remodeling enables recruitment of other cardiac transcription factors, stabilizing a CTCF-free chromatin environment productive for transcription.

We confirmed the importance of GATA4 in driving cardiac specification and maturation^52,53^. In particular, GATA4 binding to cardiac genes is fundamental for their B-to-A compartment transition, which is coupled with increased expression key for cardiomyocyte differentiation and maturation. Consistent with rat models that implicated GATA4 as a repressor of cardiac fibroblast genes^54^, GATA4 knockdown in our system led to fibroblast accumulation. In contrast to mouse studies^55^, we observed an increase in immature cardiomyocytes in a proliferative state in left ventricle organoids, suggesting a human-specific mechanism.

Much less is known about CTCF in heart development. Conditional CTCF knockout in cardiac progenitors causes severe cardiac malformation and embryonic death^56^. Interestingly, these studies reported premature arrest of cardiac development with concomitant promotion of cardiomyocyte maturation, consistent with our left ventricle single-cell data showing upregulation of electrophysiologically relevant genes such as *RYR2*. A recent mouse study found that a homozygous CTCF point mutation (R567W) severely impairs cardiac development, reducing ventricular cavity size and downregulating genes involved in sarcomere assembly and calcium handling^57^. This mutation affects the zinc finger 11, required for upstream motif recognition, and disrupts CTCF binding. In contrast, our inducible knockdown approach mimics and anticipates the physiological decrease of CTCF expression observed in the human heart and reproduced across hESC and hiPSC models. We propose that this partial and timed CTCF reduction recapitulates developmental regulation without broadly disrupting TADs and chromatin loops. Thus, whereas strong CTCF disruption is pathological, partial downregulation may represent a physiological mechanism during human cardiomyogenesis.

GATA4 and CTCF are mutated in patients with isolated CHD or in syndromes that include CHD^18,26,27,58^. The mechanism we propose may therefore provide new insight into CHD pathogenesis, though patient-specific hiPSCs will be needed to test this directly. More broadly, our work illustrates how differentiating hPSCs into human-relevant organoids that recapitulate distinct heart chambers can be leveraged to dissect CHD pathophysiology and clarify the causal links between nuclear architecture dysregulation and human disease.

Our study also reflects general challenges in separating chromatin-architectural and transcriptional mechanisms. Although the CBS edits were restricted to the core CTCF motifs, additional sequence-dependent effects cannot be fully excluded, and we therefore interpret the resulting changes as consistent with—but not exclusively due to—loss of CTCF binding. Moreover, our data cannot unambiguously separate CTCF’s architectural role from potential direct repressive activity. Both mechanisms could contribute to the observed effects, and our findings likely reflect the combined influence of CTCF on chromatin topology and gene regulation.

## Data, Code, and Material Availability

Raw NGS data for bulk and scRNA-seq have been deposited in ArrayExpress under accession numbers E-MTAB-15684, E-MTAB-15693, E-MTAB-15710, and E-MTAB-15711, while processed data is available on Zenodo (https://doi.org/10.5281/zenodo.17256127). 4C-seq data have been deposited in ENA under accession number PRJEB97970. All custom code is avail-able at GitHub (https://github.com/alessandro-bertero/Becca_et_al_2025). Computational anal-yses were conducted within containerized environments to ensure reproducibility: Hi-C (hedge-lab/hic_pipeline:image12), bulk RNA-seq (hedgelab/bulk_image:image3), single-cell RNA-seq (hedgelab/cellranger8:latest for preprocessing; hedgelabrstudio_image:iPS2seq for integration and downstream analysis), and 4C-seq (hedgelab4c:image1). All other data, code, and material used for the study is detailed in the Key Resources Table.

## Acknowledgements

We thank Giulia Savorè, Federica Sozza, Sveva Bottini, Matteo Sorge, Maddalena Arigoni, Alessia Labate, Paola Circosta, and Luca Ponzone for experimental assistance; Manuel Rosa Garrido for advice on CTCF ChIP; Estela Mancheno Juncosa for advice on cardioids; Antonio Grimaldi for MGI sequencing; and the Vienna BioCenter Core Facility for Illumina sequencing. We are also grateful to the UniTo facility staff, including Francesca Anselmi (NGS), Federica Antico (histology), Laura Conti (FACS), and Marta Gai (microscopy). This work was supported by the FEBS Excellence Award 2022 (A.B.), the Additional Ventures Single Ventricle Research Fund 2022 (A.B. and S.M.), and the Giovanni Armenise-Harvard Foundation Career Development Award 2021 (A.B.).

## Author Contributions

Conceptualization: S.Be., A.B.; Methodology: S.Be., E.M.H.; Investigation: S.Be., E.M.H., K.E.S., L.T., A.K., E.B., A.B.; Formal analysis: S.Bi.; Data curation: E.B.; Software: S.Bi., E.B.; Visualization: S.Be., S.Bi., A.B.; Resources: F.N., D.C., S.M., A.B.; Supervision: S.M., E.B., A.B.; Project administration: A.B.; Funding acquisition: A.B.; Writing – original draft: S.Be., A.B.; Writing – review & editing: all authors.

## Competing Interests

The authors declare no competing interests.

## METHODS

### Stem cell models

#### hiPSC culture

hiPSCs from an apparently healthy male donor (WTC11)^59^ and WTC11 titin-mEGFP reporter hiPSCs^60^ were cultured on Geltrex or Matrigel in Essential 8 or mTeSR Plus. Colonies were passaged as small clumps using Versene (0.5 mM EDTA in PBS) and replated in media containing 2 µM thiazovivin for 24 h. Genomic stability was verified by microarray-based karyotyping.

#### hiPSC-CM differentiation

Monolayer cardiac differentiation was performed according to a biphasic WNT signaling protocol^13^. On day –2, cells (1.5–3×10^5^ per well in 12-well plates) were seeded with 2 µM thiazovivin. On day –1, they were primed with 1 µM CHIR99021 in pluripotency medium. On day 0, differentiation was induced in RBA medium [RPMI 1640 supplemented with 0.5 mg mL^-1^ BSA and 213 µg mL^-1^ L-ascorbic acid 2-phosphate] with 5 µM CHIR99021. On day 2, 2 µM WNT-C59 was added to direct cardiac mesoderm formation. On day 4, the medium was replaced with fresh RBA, and from day 6 cells were maintained in RPMI + B27 every other day.

#### Cardiac organoids

Cardiac organoids (cardioids) were generated following the protocol of Schmidt *et al.*^30^ with minor modifications. On day –1, hiPSCs (5 × 10^4^ per well) were seeded in 24-well plates with 5 µM Y-27632. After 24 h, the medium was replaced with mesoderm (Mes) formulation. After 36 h, cells were detached with TrypLE Express, resuspended in cardiac mesoderm (CardMes) medium containing 5 µM Y-27632, and seeded into ultra-low-attachment 96-well plates (1.5 × 10^4^ cells per well), then centrifuged at 140 g for 4 min to promote aggregation. The following days (from day 2.5 onward), spheroids were fed with fresh CardMes medium without Y-27632 until day 5.5, after which they were fed with Cardio medium. All media were prepared in chemically defined medium (CDM; 50% IMDM and 50% Ham’s F12 supplemented with 0.4% BSA, 1% lipids, 0.004% monothioglycerol, and 15 µg mL^−1^ transferrin). Growth factors and small molecules were adjusted according to chamber identity. For FHF–LV, Mes contained 6 ng mL^−1^ FGF2, 5 µM LY294002, 5 ng mL^−1^ Activin A, 8 ng mL^−1^ BMP4, and 5 µM CHIR99021; CardMes contained 8 ng mL^−1^ BMP4, 1.6 ng mL^−1^ FGF2, 10 µg mL^−1^ insulin, 2 µM WNT-C59, and 50 nM retinoic acid; Cardio contained 8 ng mL^−1^ BMP4, 1.6 ng mL^−1^ FGF2, and 10 µg mL^−1^ insulin. For SHF-RV and SHF-A, Mes contained 6 ng mL^−1^ FGF2, 5 µM LY294002, 50 ng mL^−1^ Activin A, 10 ng mL^−1^ BMP4, and 6 µM CHIR99021. During CardMes (days 1.5–4.5), both RV and A were first cultured (days 1.5–2.5) with 1 μg mL^−1^ insulin (Day 1.5 only) and 10 µM SB431542; 2 µM WNT-C59 was added for RV and 5 µM XAV-939 plus 500 nM retinoic acid for A. From days 3.5–4.5, both received 10 ng mL^−1^ BMP4, 1.6 ng mL^−1^ FGF2, 10 µg mL^−1^ insulin, and 500 nM retinoic acid, maintaining 2 µM WNT-C59 for RV and 5 µM XAV-939 for A. The final Cardio (from day 5.5) contained 10 ng mL^−1^ BMP4, 1.6 ng mL^−1^ FGF2, and 10 µg mL^−1^ insulin.

### Genome editing

#### Inducible knock-down

hiPSCs were engineered with tetracycline-inducible shRNAs targeted to the *AAVS1* genomic safe harbour using an established approach^61^. Validated shRNAs from the TRC library (GATA4: TRCN0000020424; CTCF: TRCN000021849) and a scrambled control (SCR: SHC016) were cloned into pAAV-Puro_siKD by ligating annealed oligonucleotides with BglII and SalI over-hangs (Table S1). A 5’ terminal G was added to ensure efficient transcription from the H1 promoter. Cells (3 × 10^5^ per well) were seeded in 6-well plates and transfected with Gene-Juice and 2 µg total plasmid DNA [pAAV-Puro_siKD, pZFN-AAVS1_ELD, pZFN-AAVS1_KKR; equimass ratio] in Opti-MEM. Medium was refreshed after 24 h, and genome-edited cells were selected with 1 µg mL^-1^ puromycin from days 3–10, with 10 µM Y-27632 added on days 3–5. To avoid clonal bias and given the >90% on-target efficiency of this method^44^, all surviving hiPSCs were pooled for experiments. Knock-down was induced by addition of 1 µg mL^-1^ tetracycline, starting 3 days prior to differentiation, with daily medium changes.

#### CTCF binding site (CBS) editing

To identify candidate CBS within the *TTN* locus, ChIP-seq data for hESCs and hESC-derived CMs were downloaded from the ENCODE Data Portal. BigWig data were visualized on the GRCh38 genome using IGV to define six CTCF binding regions. Core motifs within each region were predicted using CTCFBSDB2.0 and confirmed with FIMO (MEME Suite). A CTCF bind-ing site score was computed with Bioconductor to estimate the degree of consensus sequence match, which served as a proxy for binding strength. To delete CBS1 and CBS6, WTC11 hiP-SCs were genome edited using single guide RNA (sgRNA)-directed Cas9 cleavage and HDR with ssODN templates. sgRNAs were designed using GuideScan2 and IDT’s CRISPR-Cas9 design checker, selecting guides with no off-targets in coding regions expressed in hiPSCs or hiPSC-CMs (CBS1: 5’-CACCGTATTCCCTCTTTGACCACTA −3’; CBS2: 5’-CACCGTATTCCC-TCTTTGACCACTA-3’). sgRNAs with BbsI overhangs were cloned as annealed double stranded oligos into pX330-U6-Chimeric_BB-CBh-hSpCas9 following standard protocols based on T4 PNK and T4 DNA ligase. The ssODN for CBS6 was synthesized on the strand opposite to the sgRNA, with 50 bp homology arms, while the ssODN for CBS1 was on the same strand, with 80 bp and 30 bp arm. Both replaced the core CTCF motif with an EcoRI site to facilitate genotyping^39^. hiPSCs were detached with Accutase and 2 × 10^6^ cells were electroporated with 2 µg pX330-gRNA plasmids (1 µg per site), 100 pmol ssODN per target, 2 µg AAVS1 SA-2A-puro-pA donor, and 2 µg each of pZFN-AAVS1_KKR and pZFN-AAVS1_ELD using the P3 Primary Cell 4D-Nucleofector Kit (CA-137 program). Cells were recovered in E8 + CEPT (50 nM chroman 1, 5 µM emricasan, 1:1000 polyamine supplement, 0.7 µM trans-ISRIB) overnight. Puromycin selection was applied to enrich HDR-competent cells^45^, followed by clonal isolation by single-cell sorting into 96-well plates. Genomic DNA was extracted, and targeted loci were amplified by PCR, digested with EcoRI to detect edited alleles, and visualized by agarose gel electrophoresis. Positive clones were validated by PCR to exclude random plasmid integra-tion (Table S2), followed by Sanger sequencing of the target regions. Heterozygous edits were resolved by TOPO-TA subcloning prior to sequencing. Selected clones were karyotyped by low-resolution microarray to confirm genomic stability.

### Functional assays

#### Flow cytometry

hiPSC-CMs were dissociated for FACS by incubating them in 0.25% trypsin for 10–15 min. The reaction was stopped with RPMI-B27 supplemented with 20% FBS, and cells were pelleted at 190 g for 5 min. After a PBS wash, cells were resuspended in PBS containing Fixable Viability Dye (eFluor 780 or 450) and incubated in the dark for 10 min. Following two washes in FACS buffer (PBS + 5% FBS), live cells were either directly analyzed for titin-mEGFP reporter activity or fixed for antibody staining. For antibody staining, cells were washed in FACS buffer twice and permeabilized during centrifugation (250 g, 5 min) in permeabilization buffer (FACS buffer + 0.75% saponin), then incubated for 50 min with either mouse anti-cardiac Troponin T (Alexa Fluor 647, 1:100) or rabbit anti-titin (targeting the Z-disc Z1Z2 region, 1:500). Titin staining was followed by a 1 h incubation with Alexa Fluor 488 goat anti-rabbit secondary antibody (5 µg mL^-1^). After washing, cells were resuspended in FACS buffer and analyzed on a FACSVerse cell analyzer, acquiring ∼30,000 events per condition. Events were gated with FlowJo v10.10.0 to exclude debris, doublets, and dead cells. For cardioid dissociation, up to 8 organoids per condition were pooled, washed twice with 0.5 mM EDTA, and incubated in 70 µL of 0.5% trypsin in EDTA at 37*^◦^*C for 10–20 min, mixing every 5 min. The reaction was stopped with CDM + 10% FBS, and cells were pelleted at 190 g for 5 min. Live titin-mEGFP recordings were performed as for hiPSC-CMs, except that FACS buffer consisted of PBS + 1% BSA.

#### Morphometry

Cardioid images were acquired daily from day 2.5 to day 7.5 of differentiation using the Incucyte SX5 Live-Cell Analysis System (Sartorius) with the spheroid analysis module and a 4× objec-tive. Size measurements were obtained by applying filters for cardioid identification, including a minimum object area of 3 × 10^4^*µ*m^2^ and a maximum eccentricity of 0.9. To exclude debris, quantification was restricted to the largest object per field.

#### Cell cycle assay

DNA synthesis was assessed using the Click-iT^TM^ EdU Alexa Fluor 647 Flow Cytometry Assay. On day 4.5, cardioids (8 per condition) were incubated with 10 µM EdU in culture medium for 2 h at 37*^◦^*C. After incubation, cardioids were dissociated as described above and stained for dead cells with Fixable Viability Dye (eFluor 780). Cells were then fixed with 50 µL Click-iT^®^ fixative (Component D) for 15 min at room temperature, washed with 1% BSA in PBS, and resuspended in Click-iT^®^ saponin-based permeabilization and wash reagent. Following 15 min permeabilization, the Click reaction cocktail was added and incubated for 30 min at room temperature in the dark. Cells were washed again with saponin-based buffer and subsequently stained with anti-titin primary and secondary antibodies (as described for flow cytometry) in the same buffer. Finally, DNA content was measured by adding FxCycle Violet Stain (1:1,000 in Click-iT^®^ saponin-based buffer) and incubating for 30 min at room temperature in the dark. Samples were analyzed on a FACSVerse cell analyzer.

#### Histology

Organoids were collected using wide-bore tips, fixed in 4% PFA at room temperature for 30 min with gentle shaking, and washed twice with PBS. Samples were incubated overnight at 4*^◦^*C in 30% sucrose/PBS, embedded in O.C.T., and cryosectioned at 12 µm using a Leica CM1950 cryostat. Sections were stored at –20*^◦^*C and thawed before staining. For staining, sections were washed in PBS for 15 min at room temperature, then permeabilized in PBST-D1 (4% goat serum, 1% Triton-X in PBS). Primary antibodies were diluted in PBST-D1 and applied within a hydrophobic barrier, incubating in the dark for 3 h at room temperature. The following primary antibodies were used: mouse anti-Ki67 (1:200), rabbit anti-α-actinin 2 (1:200), and mouse anti-pHH3 (1:50). After incubation, slides were washed in PBST2 (PBS + 0.1% Tween-20) for 15–30 min. Secondary antibodies (goat anti-rabbit Alexa Fluor 488, 1:500; goat anti-mouse Alexa Fluor 633, 1:500; goat anti-mouse Alexa Fluor 568, 1:500) were diluted in PBST2 and applied for 1 h at room temperature in the dark within a humidified chamber. After washing in PBST2, nuclei were counterstained with Hoechst 33342 (1:2000 in PBS, 5 min), then washed again. Slides were air-dried and mounted with ProLong Glass Antifade Mountant. Confocal imaging was performed using a Leica SP8 microscope with a 63X objective and the following laser settings: excitation/emission 488/520 nm (Alexa Fluor 488), 631/650 nm (Alexa Fluor 633), and 579/603 nm (Alexa Fluor 568).

#### Proliferation assay

Cell division was assessed using the CellTrace^TM^ Far Red Cell Proliferation Kit for flow cy-tometry. On day 3.5, 16 cardioids per condition were incubated with 2 µM CellTrace dye for 1 h at 37*^◦^*C in PBS containing Ca^2+^/Mg^2+^. After incubation, cardioids were washed twice with basal medium and returned to culture in complete medium. Half of the labeled cardioids were immediately dissociated and stained for live/dead discrimination (as described for flow cytome-try), and basal CellTrace fluorescence was acquired on a FACSVerse analyzer. The remaining cardioids were maintained in culture under standard conditions until day 6.5, then dissociated, stained for live/dead cells, and analyzed by flow cytometry. Cell division was quantified by calculating the median fluorescence intensity (MFI) of titin^+^ cells at day 6.5, normalized to the MFI at day 3.5. The resulting ratio for tetracycline-treated samples was then normalized to the matched untreated controls.

### Gene expression assays

#### Reverse transcription quantitative PCR (RT-qPCR)

Total RNA was extracted using the Quick-RNA MiniPrep Kit according to the manufacturer’s in-structions. cDNA synthesis was performed with the High-Capacity cDNA Reverse Transcription Kit. qPCR reactions were prepared with 10 ng cDNA, 250 nM primer mix, and either PowerUp SYBR Green 2X Master Mix or Luna^®^ Universal qPCR Master Mix. All reactions were run in technical duplicates on a QuantStudio 6 Flex Real-Time PCR System. Gene expression was normalized to the geometric mean of *RPLP0* and *PBGD* or to *HPRT1*, as indicated. Primer sequences are listed in Table S3.

#### Bulk RNA sequencing (RNA-seq)

RNA-seq for monolayer hiPSC-CMs was performed on biological duplicates using the Zymo-Seq RiboFree Total RNA Library Prep Kit with 650 ng input RNA per sample. Ribodepletion was carried out for 1 h, and libraries were amplified with 11 PCR cycles. Library quality was assessed by Qubit DNA HS and acrylamide gel electrophoresis, and sequencing was performed on an Illumina NextSeq 550 (75 bp paired-end). RNA-seq for cardiac organoids was performed on biological duplicates using the Illumina TruSeq Stranded mRNA Library Prep with 400 ng RNA input, 11 PCR cycles, and library QC by Agilent Tapestation. Sequencing was performed on an MGI T7 platform (100 bp paired-end). After demultiplexing with bcl2fastq, reads were trimmed with Trim Galore (v0.6.10) and aligned with STAR v2.7.11b using the ENSEMBL GRCh38 (release-112) genome and annotation. Gene counts were imported into R and processed with DESeq2 and the RNAseqQC package for quality control, filtering (protein-coding genes with ≥3 counts in at least two samples), and variance stabilization. Batch effects were corrected with the `removeBatchEffect` function (limma)^62^. QC included analysis of library complexity, gene detection, replicate correlation, and PCA. For expression normalization, raw counts were converted to TPM using ADImpute with gene lengths from `gtftools`. TPM val-ues were batch-corrected and used for WGCNA network analysis^63^, with GO enrichment of modules performed via `goana` and `topGO` (limma)^62^. Differential expression was conducted on raw filtered counts normalized with the `voom` function (limma), followed by contrasts testing, moderated statistics (`eBayes`), and reporting with `topTreat`. Genes were ranked by log_2_ fold change and analyzed with `gseGO` (ClusterProfiler)^64^ for biological process enrichment. For cardiac organoids, trimmed and aligned reads were analyzed as above. Heatmaps of B-to-A genes were generated from the average log_2_ fold change TPM values after batch correction, using the `pheatmap` package. Differential expression comparisons were performed separately for atrial and right ventricle genotypes. Enrichment of cardiac processes was visualized with `gseaplot` (enrichplot). Rank-rank hypergeometric overlap (RRHO) analysis was performed with RedRibbon, comparing log_2_ fold changes of CTCF (TET vs. no TET) and GATA4 (TET vs. no TET) in atrial and ventricular samples. RRHO maps were visualized with the `ggRedRibbon` function.

#### Single-cell RNA-seq (scRNA-seq)

FHF-LV cardioids at days 4.5 and 7.5 were analyzed in biological duplicates by microfluidics-based scRNA-seq (10X Genomics). Sixteen cardioids per condition were pooled, dissociated with 0.5% trypsin/0.5 mM EDTA, quenched with RPMI-B27 + 20% FBS, centrifuged (200 g, 5 min, 4*^◦^*C), and resuspended in PBS/1% BSA. Cells were counted and labeled with barcoded lipid-conjugated cell multiplexing oligos (CMOs; 0.5 million cells/condition, 100 µL, 5 min, RT), washed, stained with Fixable Viability Dye, and live-sorted on a Sony SH800S (100 µm chip, targeted mode, single-cell 3-drop). Controls (2.5 × 10^5^ cells each) and tetracycline-treated samples (7.5 × 10^5^ cells) were pooled, and 5.0 × 10^5^ total cells were loaded per channel on the Chromium Next GEM Chip G. Libraries were prepared with the Chromium Next GEM Single Cell 3’ Reagent Kit v3.1 and sequenced on an Illumina NovaSeq 6000 (S2 100 cycles, 18+90) to a depth of ∼4 billion reads (∼18,000 singlets/condition, ∼50,000 reads/GEX, ∼7,500 reads/CMO per cell). A similar workflow was applied to pooled FHF-LV, SHF-LV, and SHF-A cardioids (3.0 × 10^5^ cells total), sequenced on an Illumina NextSeq 1000 (P2 100 cycles). Raw fastq files were processed with Cell Ranger v8.0.1 (multi, aggr without normalization) against GRCh38. Count matrices were imported with `Read10X` and analyzed with Seurat. Filtering criteria included ≥200 expressed genes per cell, genes expressed in ≥3 cells, mitochondrial content ≤15%, and removal of outliers (lowest 5% / highest 1% by UMI or gene counts). Data were normalized with `NormalizeData` (log, scale factor 10,000), scaled with `ScaleData`, and scored for cell cycle phases using `CellCycleScoring`. Filtered matrices were imported into Monocle3 for trajectory analysis. Dimensionality reduction was performed with PCA and UMAP; cluster-ing with `cluster_cells` (partition q-value 0.05); and pseudotime inference with `learn_graph` and `order_cells`, rooting at the branch enriched for pluripotency markers. Cluster stability was assessed by resampling (10 iterations, 1,000 cells removed each). Marker genes were identi-fied with `top_markers`, and GO enrichment was performed with `goana` and `topGO`. Differential expression was tested at the cluster level (`fit_models`, Poisson; GSEA with `gseGO`) and along pseudotime (`graph_test`, q < 0.05). Cardiomyocyte clusters were further sub-analyzed by pairwise comparisons (`fit_models`, negative binomial), followed by enrichment with `gseGO` and `gseKEGG`. For condition-specific comparisons, untreated controls (all genotypes pooled) were compared to each tetracycline-treated genotype using Fisher’s exact test with multiple-testing correction (`p.adjust`). Finally, the differential abundance of cells per cluster was assessed using MiloR. kNN graphs were built with `buildGraph` (k = 30, 30 dimensions), and overlapping neighborhoods were generated with `makeNhoods`. Neighborhood distances were computed with `calcNhoodDistance` (d = 30), and differential abundance was tested with `testNhoods`, using shRNA-matched CTR as the reference condition. Results were summarized at the cluster level by computing the median logFC, the fraction of significant neighborhoods, direction consistency, and the minimum spatialFDR. Clusters were considered significant when minSpatialFDR ≤ 0.05 with median logFC > 0.5 or < –0.5 and frac_sig > 0.75. Cluster ID transferring was performed by importing GSE263193 scRNAseq counts into Seurat, filtering for genes expressed in both datasets, performing log_2_ normalization and data processing with the default Seurat pipeline. Transferring anchors were identified with the `FindTransferAnchors` function and transferred with the `TransferData` function.

### Chromatin and nuclear architecture assays

#### Chromatin immunoprecipitation (ChIP)-qPCR

Chromatin immunoprecipitation (ChIP) for CTCF was performed with minor modifications of a published protocol^65^. Cells were crosslinked in 1% formaldehyde for 15 min at room temperature with gentle agitation, and the reaction was quenched with 125 mM glycine for 10 min. All subsequent steps were performed on ice unless otherwise indicated. After washing with cold PBS, cells were scraped, pelleted (250 g, 5 min, 4*^◦^*C), and lysed in cell lysis buffer (10 mM Tris-HCl pH 8.0, 10 mM NaCl, 0.2% NP-40) supplemented with protease inhibitors. Nuclei were isolated by centrifugation (600 g, 5 min), then lysed in nuclear lysis buffer (50 mM Tris-HCl pH 8.0, 10 mM EDTA, 1% SDS). Chromatin was diluted in buffer (20 mM Tris-HCl pH 8.0, 2 mM EDTA, 150 mM NaCl, 0.01% SDS, 1% Triton X-100) and sheared with a Bioruptor Pico (7 cycles of 30 s ON/30 s OFF). The lysate was clarified (16,000 g, 10 min, 4*^◦^*C), and an aliquot was saved as input. The remaining solution was incubated overnight at 4*^◦^*C with a mix of 2 µL rabbit anti-CTCF and 8 µL mouse anti-CTCF antibodies, or with matched rabbit IgG and mouse IgG1*κ* isotype controls. Immunocomplexes were captured for 1 h at room temperature with Protein A/G magnetic beads, which had been pre-washed in wash buffer. Beads were sequentially washed twice with buffer I (20 mM Tris-HCl pH 8.0, 2 mM EDTA, 50 mM NaCl, 0.1% SDS, 1% Triton X-100), once with buffer II (10 mM Tris-HCl pH 8.0, 1 mM EDTA, 0.25 M LiCl, 1% NP-40, 1% sodium deoxycholate), and twice with TE (10 mM Tris-HCl pH 8.0, 1 mM EDTA). DNA was eluted twice in 100 mM sodium bicarbonate, 1% SDS, for 15 min each at room temperature. Eluates and input were treated in parallel with 1 µg RNase A and 300 mM NaCl, then incubated 5 h at 65*^◦^*C to degrade RNA and reverse crosslinks. Samples were then treated overnight with 60 µg Proteinase K at 45*^◦^*C and purified with the ChIP DNA Clean & Concentrator kit. qPCR was performed on target regions using 4 µL purified DNA, 5 µL 2X Luna Universal qPCR Master Mix, and 2.5 µM primer mix (Table S4). Ct values were normal-ized to inputs and then to the average Ct of four negative control regions lacking CTCF binding and compared to two constitutively-bound positive controls on the same chromosome (hg38 chr2:176140583-176140677 and chr2:176140649-176140792).

#### Circularized Chromosome Conformation Capture with sequencing (4C-seq)

4C was performed following an established protocol^66^ with minor modifications. hiPSCs were detached, pelleted (100 g, 5 min), and resuspended at 2 × 10^6^ cells/mL in isolation buffer (10% FBS in PBS). Fixation was carried out by adding an equal volume of 4% formaldehyde in isolation buffer and incubating for 10 min at room temperature with shaking, followed by quenching with 0.13 M glycine for 5 min. Cells were pelleted (500 g, 5 min, 4*^◦^*C), washed in PBS, and lysed in ice-cold lysis buffer (50 mM Tris-HCl pH 8.0, 0.5% NP-40, 1% Triton X-100, 150 mM NaCl, 5 mM EDTA) supplemented with protease inhibitors. Nuclei were pelleted (500 g, 5 min, 4*^◦^*C) and resuspended in 1.2× RE1 buffer with 0.3% SDS, incubated for 1 h at 37*^◦^*C with shaking, then treated with 2.5% Triton X-100 for 1 h under the same conditions. Digestion was performed with 100 U DpnII for 3 h at 37*^◦^*C, followed by an additional 100 U overnight. Digestion efficiency was verified on agarose gel. After heat inactivation, samples were ligated overnight at 16*^◦^*C in 2 mL reactions containing 50 U T4 ligase. Crosslinks were reversed with 75 µg/mL proteinase K, and DNA was purified by isopropanol precipitation and ethanol washes. DNA was eluted in nuclease-free water and subjected to a second digestion with Csp6I (overnight at 37*^◦^*C, 20 min at 65*^◦^*C, then 4*^◦^*C), followed by a second ligation (25 µg DNA template in 5 mL reaction, 50 U ligase, overnight at 16*^◦^*C). DNA was purified again and quantified. PCR was performed in two steps: (1) first-round amplification with LongAmp Taq polymerase (16 cycles, annealing 50*^◦^*C) using primers with partial Illumina overhangs (reading primer: 5’-TACACGACGCTCTTCCGATCTNNNNNAACCAGCTTAAATTGATC-3’; non-reading primer: 5’-ACTGGAGTTCAGACGTGTGCTCTTCCGATCTTTCAGAACAGAATAGTCATG-3’); (2) second-round amplification with NEBNext High-Fidelity PCR Mix and Illumina dual indexes. PCR products were size-selected with SPRI beads (0.8× ratio). The viewpoint was defined at *chr2:178,805,901–178,806,592*, and 800 ng DNA was used for amplification. Libraries were sequenced on an Illumina NextSeq 1000 (100 cycles kit, 58 cycles read 1). Bioinfor-matic analysis followed the 4C-seq pipeline^66^ within a Docker container (hedgelab/4c:image1). Primer sequences, restriction enzymes (DpnII: GATC; Csp6I: GTAC), and genome reference (hg38) were specified in the viewpoint configuration file. Reads were demultiplexed by primer, trimmed at the 5’-end (DpnII site), mapped to hg38 with bowtie2, and filtered for overlaps with the in silico digested genome. After read counting and smoothing, wig files were converted to bed format (wig2bed, bedops) and imported into R. Batch correction was performed with `removeBatchEffect` (limma), normalizing z-scores across replicates. Data were restricted to bins overlapping *TTN*, excluding the viewpoint. Area-under-curve (AUC) values between and under CTCF ChIP peaks (ENCFF332TNJ) were computed with `AUC` (DescTools). Difference curves were generated by reciprocal subtraction between conditions with the pyGenomeTracks module^67^, and the difference in each region was tested with a one sample t-test (mu = 0). Comparison with virtual 4C tracks (RUES2 hESCs, hESC-MES, hESC-CPs, and hESC-CMs Hi-C data) was performered by binning the 4C tracks at 40kb, filtering for bins on chromosome2, rescaling each track on the library size, log_2_ normalizing and fitting on a 20-degree polynomial curve with the `lm` function. For each pair of conditions it was evaluated the Pearson correlation of the polynomial coefficients of the derivatives with the `rcorr` function.

#### Immunofluorescence paired to DNA fluorescence *in situ* hybridization (Immuno-FISH)

ImmunoFISH following a published protocol^13^ with minor modifications. The *TTN* locus-spanning CH17-275G10 BAC (BACPAC Resources Center) was purified with the Qiagen Plasmid Midi Kit, with modified centrifugation steps to preserve large BAC DNA. The product was validated by PCR using two primer pairs amplifying the expected region (5’-AGTAAGGCGGGTGGCTTTAT-3’ – 5’-CCACATCACAACCGTGAAAG-3’, T_a_ 56*^◦^*C; 5’-AGGCTGGCTGAAAAATAGGTC-3’ – 5’-TAATACAACGCATTGGCCCTC-3’, T_a_ 57*^◦^*C). Probes were labeled by nick translation with Alexa Fluor 488-dUTP (4 h at 15*^◦^*C, inactivation at 70*^◦^*C, 10 min), purified, and precipitated with salmon sperm DNA, human Cot-1 DNA, sodium acetate, and ethanol overnight at –20*^◦^*C. The pellet was resuspended in hybridization buffer (50% formamide, 2× SSC, 10% dextran sulfate). hiPSC-derived cardiomyocytes (day 24) underwent a 45 min heat shock at 42*^◦^*C in RPMI + HEPES, were returned to 37*^◦^*C overnight, and replated (40,000 cells/slide) on geltrex-coated HistoBond slides. After 4 h, cells were fixed in 4% PFA (10 min), quenched with 0.1 M Tris-HCl pH 7.4 (10 min), and cryoprotected in 20% glycerol (20 min) followed by 50% glycerol overnight at –20*^◦^*C. Slides were permeabilized and blocked in 0.1% Triton X-100 + 1% goat serum in PBS (1 h), incubated overnight at 4*^◦^*C with rabbit anti-Lamin B1 (1:350) and mouse anti-α-actinin (1:100), and washed. Secondary antibodies (goat anti-rabbit Alexa Fluor 405, 1:500; goat anti-mouse Alexa Fluor 633, 1:500) were applied for 1 h at room temperature. Post-fixation was performed with 3% PFA in PBS (10 min). Slides were treated with 0.1 M HCl + 0.7% Triton X-100 (10 min, 4*^◦^*C), washed in SSC, and DNA was denatured in 50% formamide/2× SSC (30 min, 80*^◦^*C). Denatured probes (10 µL/slide) were applied under coverslips sealed with rubber cement, and hybridized overnight at 42*^◦^*C. Slides were washed in 50% formamide/2× SSC (3×, 42*^◦^*C), then in 2× SSC at 42*^◦^*C and room temperature. Slides were mounted in ProLong Diamond antifade, dried 2 h, sealed, and imaged within a few days. Images were acquired on a Zeiss Elyra 7 microscope (63×, NA 1.4, HiLo mode). Z-stacks encompassed the Lamin B1 signal. At least 50 cardiomyocytes per condition were imaged. Analysis was performed in Imaris v10.2: nuclei were segmented with a surface filter (0.5 µm), Lamin B1 and FISH background removed with size filters (9 µm and 1 µm, respectively), and only α-actinin^+^ cells retained. Distances between FISH spot centers and nuclear boundaries were computed with the distance transformation function, normalized to the average cube root of the nuclear volume, and batch processed with a custom Python script. Non-diploid nuclei (≠2 spots) were excluded. The number of cardiomyocytes analyzed per condition is reported in the figures.

#### Bisulfite pyrosequencing

Bisulfite conversion of genomic DNA was performed using the Zymo Methylation-Lightning Kit following the manufacturer’s instructions. Bisulfite-treated DNA was amplified with the Pyro-Mark PCR Kit, and PCR products were sequenced on a PyroMark Q48 Autoprep using Pyro-Mark Q48 Advanced CpG Reagents. DNA methylation levels were quantified with the PyroMark Q48 Autoprep Software. Primers were designed with the PyroMark Assay Design Software 2.0. The forward primer was 5’-GAATTTGGTTTTTTATTATTGGTAAGATTG-3’, the reverse primer (biotinylated) was 5’-CTCCAAATAAATTACTCCATAAATCATCTA-3’, and the sequenc-ing primer was 5’-GTTTTTTATTATTGGTAAGATTGTA-3’.

### Secondary data integration and analysis

#### Hi-C data analysis

DNase Hi-C datasets for RUES2 hESCs, hESC-MES, hESC-CP and hESC-CMs were down-loaded from the 4DN Data Portal. Biological replicates were kept separate, while technical replicates were concatenated using the `cat` function. Reads were aligned with HiC-Pro v3.1.0^68^ to the hg38 genome, indexed with `bowtie2-build` (UCSC FASTA and chromosome sizes files, excluding chrX, chrY, and chrM). Default parameters were used, except `GENOME_FRAGMENT` and `LIGATION_SITE`, which were left empty as required for DNase Hi-C. Differential compartment analysis was performed with the dcHiC pipeline^69^ using ICE-normalized matrices binned at 150 kb. PCA was first computed per chromosome (`pcatype-cis`), the optimal component was se-lected (`pcatype-select`), and differential analysis was conducted (`pcatype-analyze`) to gener-ate IGV tracks (`pcatype-viz`). Raw HiC-Pro matrices at 40 kb resolution were converted to `h5` format with HiCExplorer v3.4.1^70^ (`hicConvertFormat.py`). Biological replicates were summed (`hicSumMatrices.py`), filtered and ICE-normalized (`hicCorrectMatrix.py`), then converted to `mcool` format and imported into R with `HiCExperiment`. Virtual 4C was performed with `virtual4C` (HiContacts package)^71^ using the *TTN* promoter (hg38 chr2:178,806,793–178,808,363) as viewpoint. BedGraph outputs were visualized in IGV and final tracks plotted with pyGenome-Tracks. For H9 hESC-CM differentiation time-course Hi-C, raw `hic` matrices were downloaded from GEO and converted to HiC-Pro–like sparse matrices with `preprocess.py` (dcHiC envi-ronment). Chromosomes X, Y, and M were excluded, and resolution was set to 50 and 100 kb. ICE normalization was performed with HiC-Pro v3.1.0 (`step` `=` `ice_norm`, default settings). Differential compartments were identified with dcHiC using 100 kb bins. Virtual 4C was per-formed at 50 kb resolution as above, with the viewpoint lifted from hg19 to hg38 (UCSC LiftOver; hg19 coordinates: chr2:179,671,520–179,673,090). For WTC11 hESC-CM, normalized `mcool` matrices were downloaded from the 4DN DataPortal^14^ and converted to HiC-Pro–like sparse matrices with `preprocess.py` at 50kb and 100kb resolutions. Differential compartment analysis (100kb) and virtual4C analysis (50kb) were performed as described above.

#### ChIP-seq data analysis

Publicly available GATA4 ChIP-seq datasets in hESCs and CMs (aligned to hg38) were pro-cessed as previously described^13^. CTCF ChIP-seq for hESC (ENCSR000AMF) and CM (ENCSR713SXF), and RAD21 ChIP-seq for hESC (ENCSR000ECE) were downloaded from ENCODE in `bed` (pseudo-replicated IDR-thresholded peaks) and `BigWig` (fold-change over control) formats, us-ing the ENCODE4 v1.5.1 GRCh38 annotation. Each `BigWig` file was visualized together with its corresponding `bed` file using pyGenomeTracks, following manual customization of tracks generated with the `make_tracks_file` function. The bin size was set to 10 bp in the region of interest to enable high-resolution visualization. Genomic coordinates of CTCF discrete peaks identified at each developmental stage were compared, and a peak was classified as conserved if an overlap was detected.

#### ATAC-seq data analysis

Publicly available ATAC-seq datasets were downloaded from GEO in `BigWig` format. For each condition, the signal was averaged across the two biological replicates and visualized with pyGenomeTracks. Binning was set to 10 bp within the region of interest.

#### Single-cell data integration

Single-cell transcriptomic datasets were obtained from human preimplantation embryos (day 3–7, ArrayExpress), human gastrula (day 16, ArrayExpress), and human fetal hearts (week 5–25, GEO). Raw counts were downloaded and only undifferentiated and mesendoderm-committed cells (for early datasets) were retained. A unified count matrix was generated using the intersec-tion of gene names across datasets. Cells with fewer than 200 expressed genes were excluded, and the filtered matrix was imported into a Monocle3 `cell_dataset` object. Data were nor-malized with `preprocess_cds` (PCA, log normalization, 50 dimensions), then batch-corrected using `align_cds`. Dimensionality reduction was performed with UMAP (`reduce_dimension`, `umap.min_dist = 1`), followed by clustering (`cluster_cells`), trajectory inference (`learn_graph`), and pseudotime ordering (`order_cells`), rooting at the branch containing day 3 preimplantation cells. Because trajectories showed reduced linearity, data were further subdivided by cham-ber identity (RA, RV, LA, LV) based on the fetal heart dataset classification. Each subset was reprocessed through the same Monocle3 pipeline. UMAPs and pseudotime trajectories were vi-sualized with `plot_cells` and `plot_genes_in_pseudotime` to examine chamber-specific gene expression dynamics.

#### Cardiac B-to-A and A-to-B gene analysis

Cardiac B-to-A genes were identified from H9 cell Hi-C compartmentalization data, which pro-vided the deepest sequencing coverage and the widest time-course (day 0 to day 80 of car-diomyocyte differentiation). 100 kb bins were classified based on the compartment assigned with dcHiC (as described above) at day 0 and at day 80 in A-to-A, A-to-B, B-to-B, and B-to-A. Genes belonging to each bin were then defined as those with transcription start sites (TSS) located within these bins. To assess expression dynamics, bulk RNA-seq data associated with the H9 Hi-C dataset were analyzed. RPKM-normalized counts were downloaded from GEO and converted to TPM values. TPM counts were batch-corrected across replicates using the `removeBatchEffect` function (limma), and differential gene expression analysis from day 0 to day 80 was performed using the limma pipeline described above. Significantly differentially expressed genes were classified based on a q-value threshold of 0.05 and an absolute log_2_ fold change of 2. These selected genes were used to annotate the bins in the 4 categories of compartment changes with the number of observed up and downregulated genes. These values were compared to the expected probability of each category computed based on the gene density and the total percentage of disregulated genes in each direction using a chi-squared test with the “chisq.test” R function (”stats” package). Individual p-values were computed as a 2-sided z-test on the distribution of the residues. Gene set enrichment analysis (GSEA) was performed with `fgsea`^72^, using the list of up- and down-regulated genes in each compartment category as pathways, and genes ranked by log_2_ fold change from differential expression analysis (tetracycline vs. isogenic control) in both bulk RNA-seq datasets. Enrichment curves were plotted with `plotEnrichment` for each comparison.

### Statistics

Statistical analyses were performed using GraphPad Prism v10. The number of biological replicates, statistical test applied, and exact p-values are reported in the figure legends. Unless otherwise specified, data are presented as mean *±* SEM.

### Key resources table

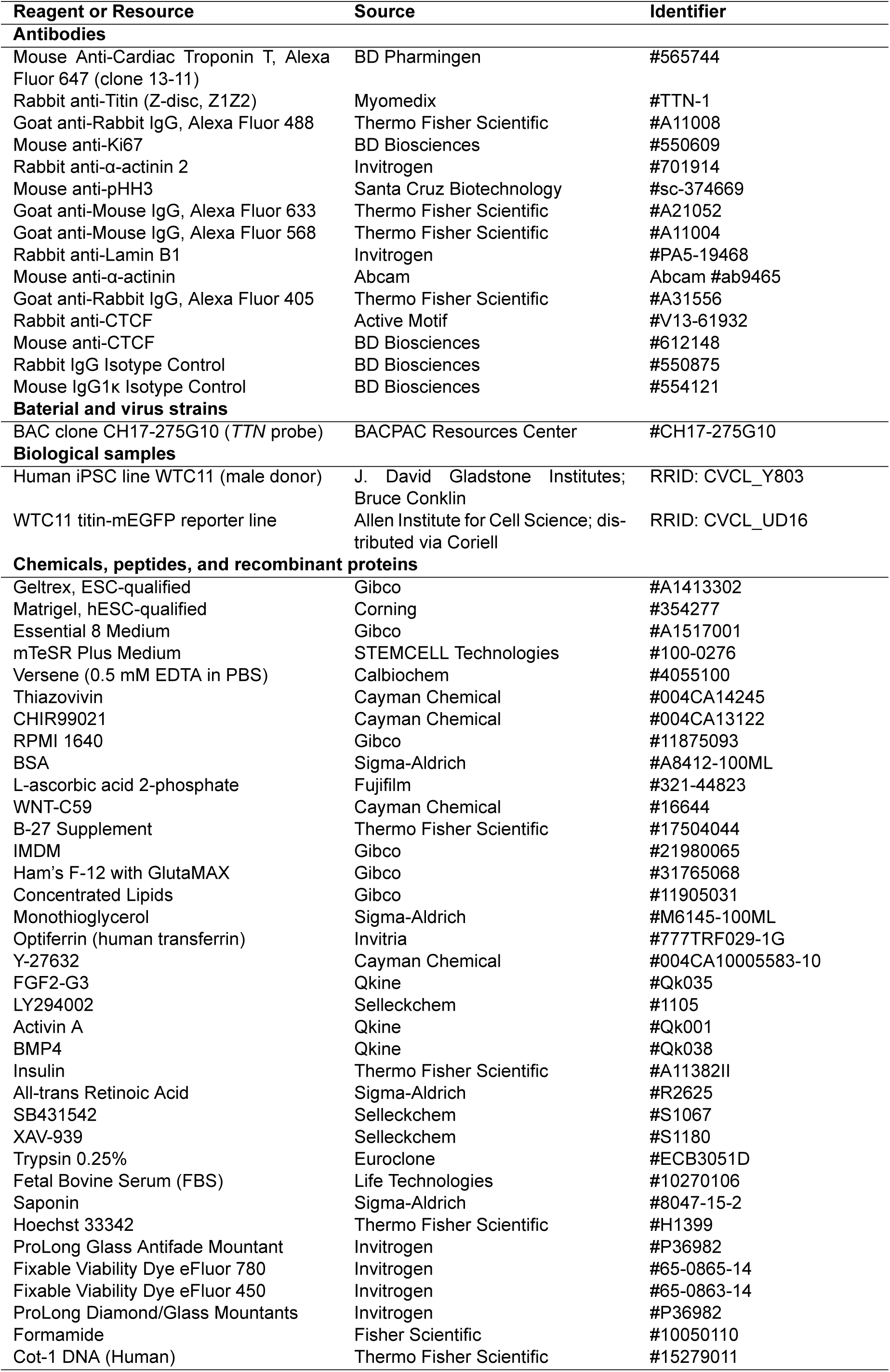
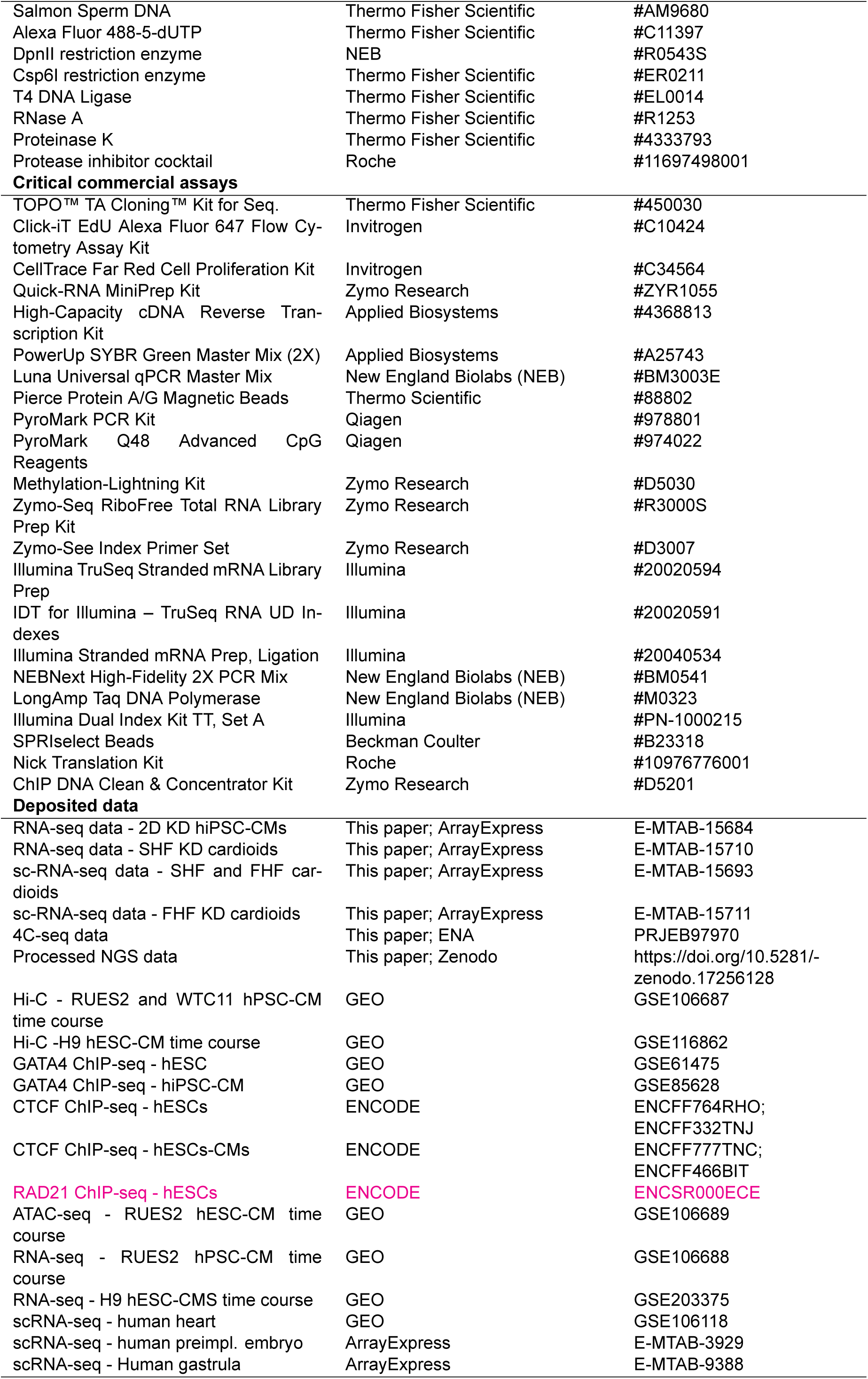
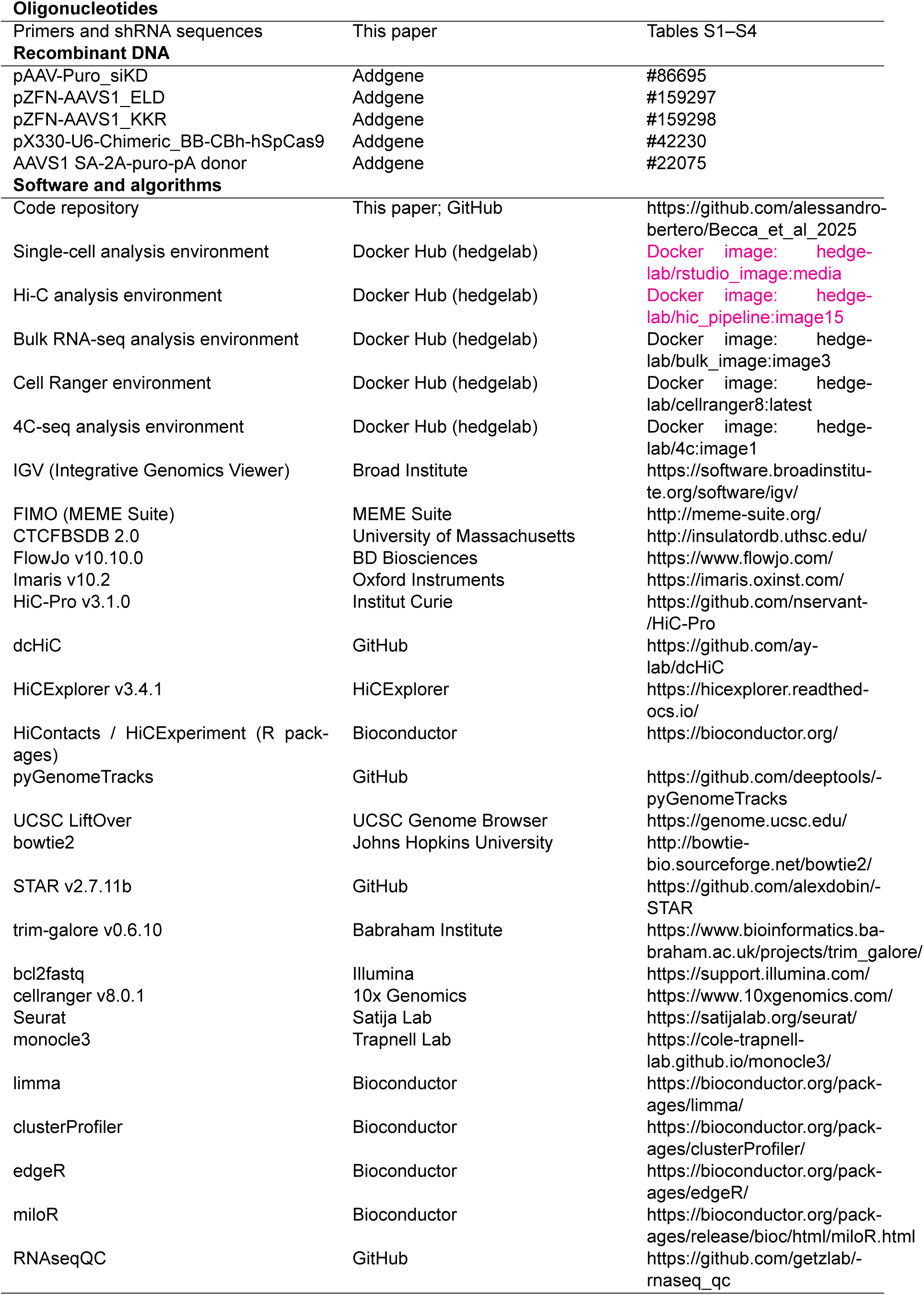
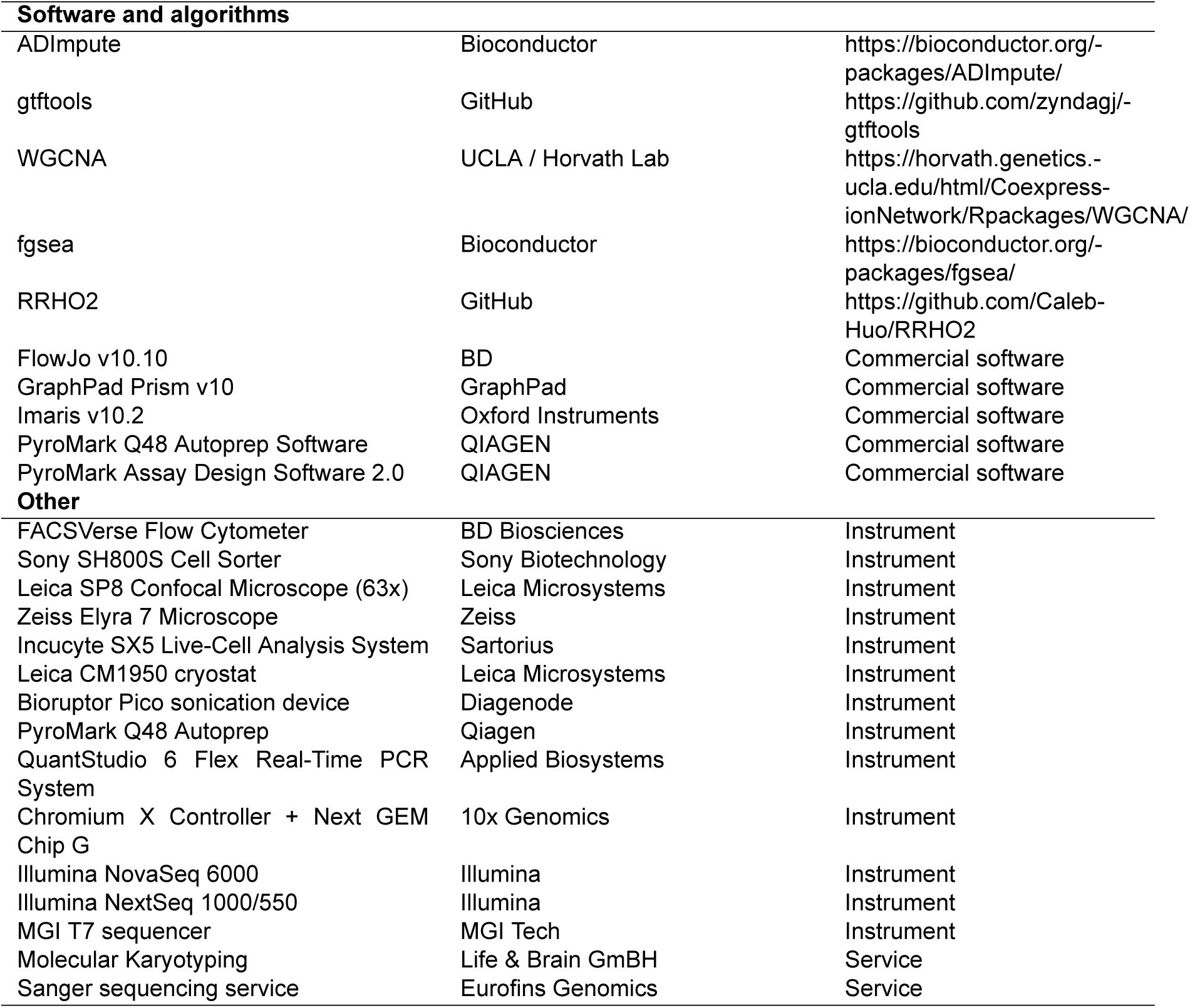

## Notes

https://zenodo.org/records/17256127

https://github.com/alessandro-bertero/Becca_et_al_2025

### Competing Interest Statement

The authors have declared no competing interest.

### Summary of Updates

This is a preliminary revision in response to the peer review performed at Review Commons. All changes are described in detail in our point-by-point responses to the reviewer's comments.

