## Supplemental Information for "Opposing CTCF and GATA4 activities set the pace of chromatin topology remodeling during cardiomyogenesis"

### Supplemental Items

- Figure S1. Reciprocal dynamics of the *TTN* locus regulators GATA4 and CTCF
- Figure S2. Inducible knockdown of GATA4 or CTCF during hiPSC-CM differentiation
- Figure S3. Precise disruption of CTCF binding sites at *TTN*
- Figure S4. Single-cell RNA-seq characterization of FHF and SHF cardioids
- Figure S5. Inducible knockdown during SHF cardioid differentiation
- Figure S6. Single-cell RNA-seq of differentiating inducible knockdown LV cardioids
- Figure S7. Proliferation dynamics in inducible knockdown LV cardioids
- Table S1. Oligonucleotides for molecular cloning
- Table S2. Oligonucleotides for genotyping
- Table S3. Oligonucleotides for RT-qPCR
- Table S4. Oligonucleotides for ChIP-qPCR

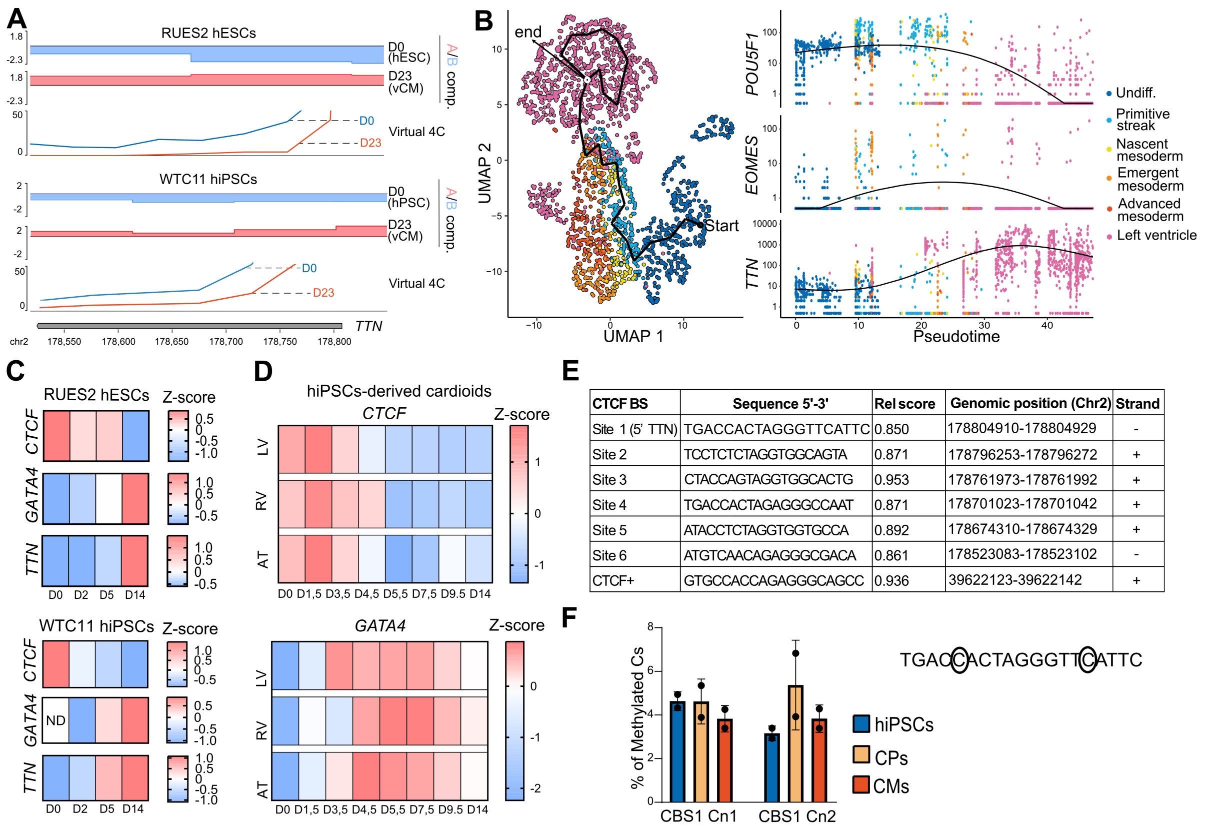


| CTCF BS | Sequence 5'-3' | Rel score | Genomic position (Chr2) | Strand |
|---|---|---|---|---|
| Site 1 (5' TTN) | TGACCACTAGGGTTCATTC | 0.850 | 178804910-178804929 | - |
| Site 2 | TCCTCTCTAGGTGGCAGTA | 0.871 | 178796253-178796272 | + |
| Site 3 | CTACCAGTAGGTGGCACTG | 0.953 | 178761973-178761992 | + |
| Site 4 | TGACCACTAGAGGGCCAAT | 0.871 | 178701023-178701042 | + |
| Site 5 | ATACCTCTAGGTGGTGCCA | 0.892 | 178674310-178674329 | + |
| Site 6 | ATGTCAACAGAGGGCGACA | 0.861 | 178523083-178523102 | - |
| CTCF+ | GTGCCACCAGAGGGCAGCC | 0.936 | 39622123-39622142 | + |

#### Figure S1. Reciprocal dynamics of the *TTN* locus regulators GATA4 and CTCF

(A) Global chromatin topology remodelling at *TTN* during hPSC-CM differentiation, assessed by Hi-C in RUES2 hESCs and WTC11 hiPSCs (data plotted from quantile normalized average of n = 2 differentiations)[13]. Virtual 4C extracted from Hi-C data shows *TTN* promoter interactions. vCM: ventricular cardiomyocytes. Data plotted from merged n = 2 differentiations for RUES2 hESCs or n = 1 differentiation for WTC11 hiPSCs

(B) Uniform manifold approximation and projection (UMAP) of single-cell RNA-seq data of early human embryos[34–36], with selected markers genes for pluripotency (*POU5F1/OCT4*), primitive streak (*EOMES*), and cardiomyocytes (*TTN*). Normalized read counts are shown. Cells are ranked developmentally based on pseudotime, with color coding indicating cell type.

(C) Bulk RNA-seq time course during cardiac differentiation of RUES2 hESCs[13] and RT-qPCR of WTC11 hiPSCs. Average z-scores from n = 2 differentiation for RUES2 hESCs and n = 3 differentiations for WTC11 hiPSCs. ND: non-detected.

(D) Bulk RNA-seq time course during chamber-specific cardiac organoid differentiation of WTC11 hiPSCs (GSE239891). n = 1 differentiation. LV: left ventricle; RV: right ventricle; AT: atria.

(E) Table of *TTN* CTCF binding site (CBS) core sequences and a positive control site, showing genomic locations and conservation scores.

(F) DNA methylation levels of *TTN* CBS1 cytosines measured by pyrosequencing. n = 2 differentiations. CP: cardiac progenitors.

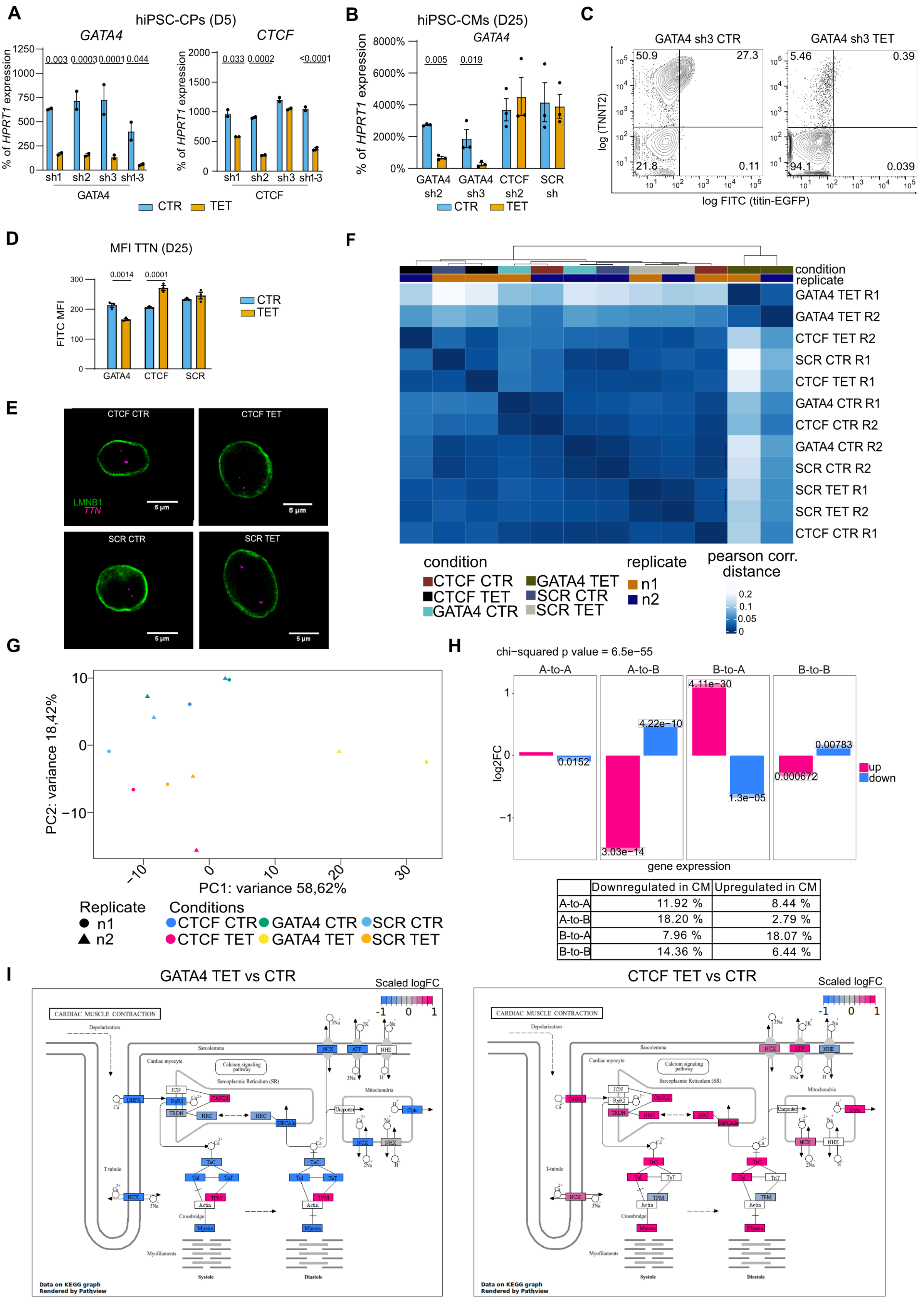


| | Downregulated in CM | Upregulated in CM |
|---|---|---|
| A-to-A | 11.92 % | 8.44 % |
| A-to-B | 18.20 % | 2.79 % |
| B-to-A | 7.96 % | 18.07 % |
| B-to-B | 14.36 % | 6.44 % |

**Figure S2. Inducible knockdown of GATA4 or CTCF during hiPSC-CM differentiation**

Legend on the next page

#### Figure S2. Inducible knockdown of GATA4 or CTCF during hiPSC-CM differentiation (continued)

(A) RT-qPCR validation of knockdown efficiency using three different inducible shRNAs and their combination in hiPSC-derived CPs (day 5). n = 3 wells, p-values by two-way ANOVA with Holm-Sidak corrected pairwise multiple comparisons. CTR: no tetracycline control; TET: tetracycline.

(B) RT-qPCR validation of *GATA4* knockdown in hiPSC-CMs (day 25), as reported in Figure 2A, here including shRNA 3. n = 3 differentiations, p-values by two-way RM ANOVA with Holm-Sidak corrected pairwise multiple comparisons.

(C) Representative flow cytometry plots showing cardiac differentiation efficiency for the *GATA4* shRNA 3 condition (compare to Figure 2C).

(D) Titin-EGFP median fluorescence intensity (MFI) in day 25 hiPSC-CMs. n = 3 differentiations, p-values by two-way RM ANOVA with Holm–Sidak corrected multiple comparisons.

(E) Representative images of immunoFISH used to quantify the distance of the *TTN* locus from the nuclear lamina (LMNB1) in inducible knockdown hiPSC-CMs (refer to Figure 2D). Individual sections of Z-stacks used for 3D volumetric reconstruction.

(F) Pearson correlation of bulk RNA-seq data considering the 1,000 most variable genes in inducible knockdown hiPSC-CMs. n = 2 differentiations (R1 and R2).

(G) Principal component analysis (PCA) of the 1,000 most variable genes from the RNA-seq shown in panel E.

(H) Chi-squared test of differential compartment changes in H9 hESCs from day 0 to day 80 of cardiac differentiations (refer to Figure 1A), correlating nuclear compartment changes with gene expression changes (p-value on the barchart and table on the bottom).

(I) Pathview visualization of the KEGG muscle contraction pathway (hsa04260) from the RNA-seq data shown in panel E for genes with p-adjusted < 0.05.

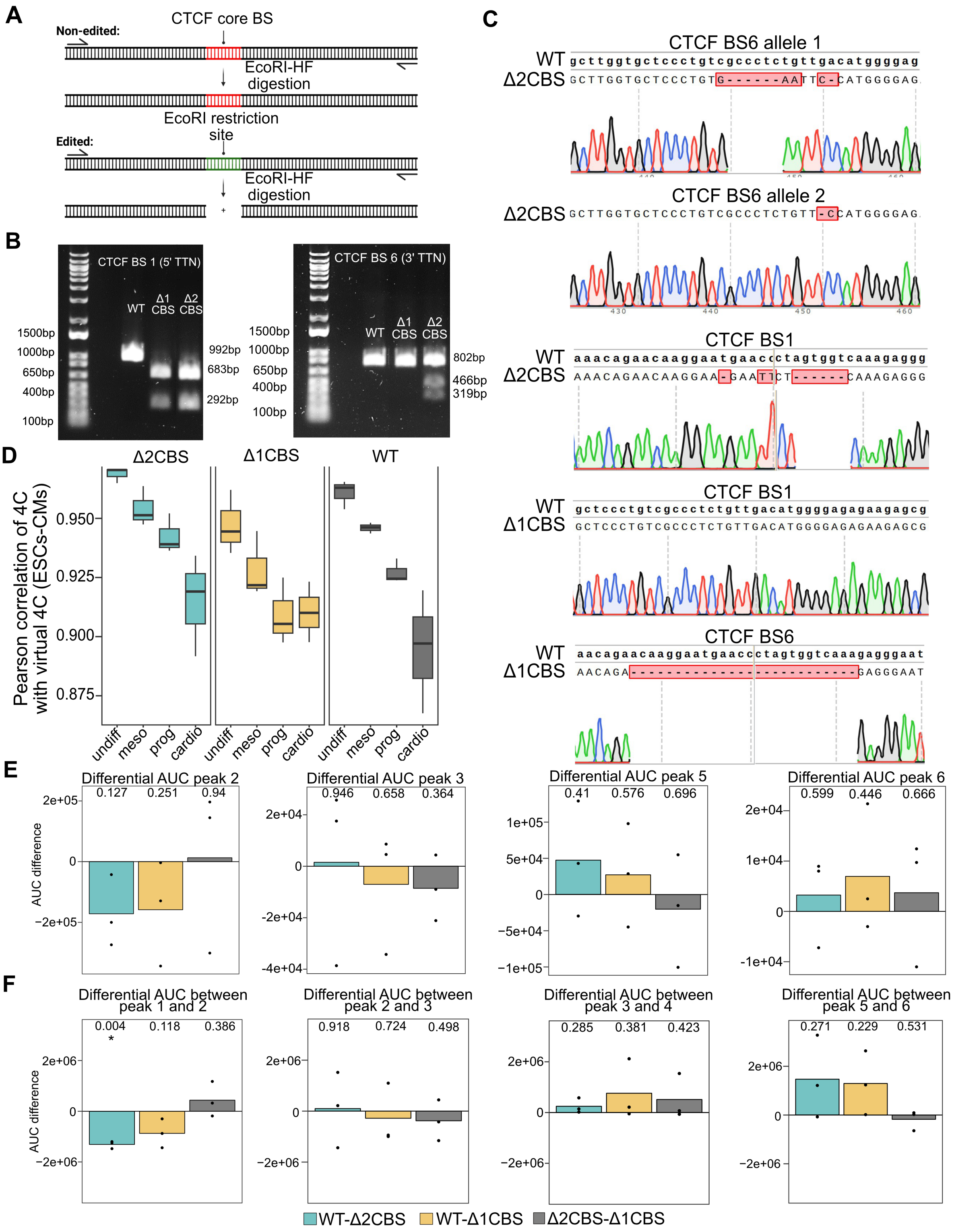


#### Figure S3. Precise disruption of CTCF binding sites at *TTN*

Legend on the next page

#### Figure S3. Precise disruption of CTCF binding sites at *TTN* (continued)

(A) Schematic of the genotyping strategy for edited CTCF binding sites (CBSs; refer to Figure 3A).

(B) Genotyping results for selected genome-edited hiPSC clones. PCR amplicons spanning the targeted regions were digested with EcoRI to test for replacement of the core CTCF motif with the restriction site. Δ1CBS: homozygous deletion of CBS1; Δ2CBS: homozygous deletion of CBS1 plus compound heterozygous disruption of CBS6; WT: wild-type.

(C) Sanger sequencing of TOPO-cloned PCR amplicons of CBS6 from a putative heterozygous-edited clone. The second allele carried a 2-bp deletion within the CTCF motif. Representative sequences of homozygous edits are also shown.

(D) Correlation of 4C-seq interaction profiles of the *TTN* promoter in WT and edited hiPSCs, compared with virtual 4C derived from Hi-C during cardiac differentiation of RUES2 hESCs[13].

(E) Differential 4C-seq AUC between pairs of conditions for the indicated CBSs. n = 3 cultures, p-values by one-sample t-test against $\mu = 0$.

(F) As in panel E, but for the indicated genomic intervals.

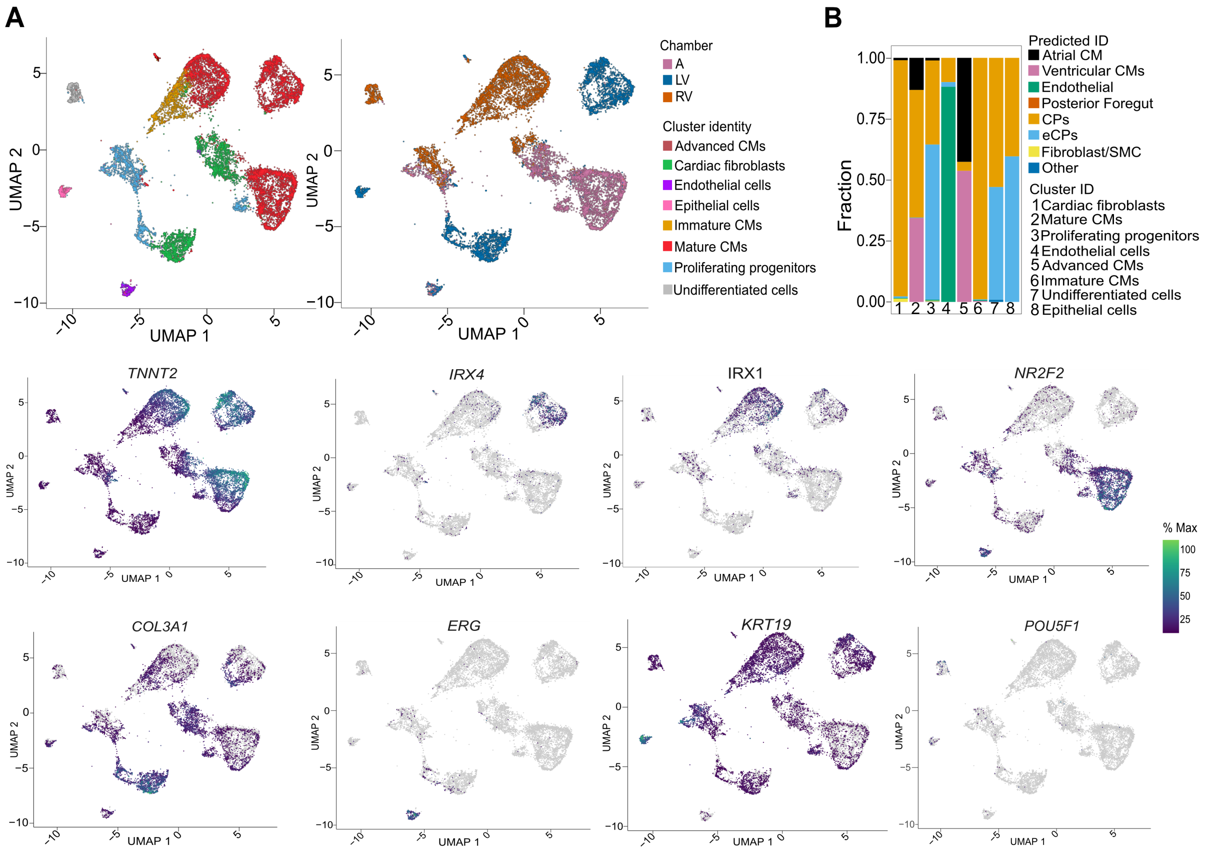


**Figure S4. Single-cell RNA-seq characterization of FHF and SHF cardioids**

(A) UMAP representation of scRNA-seq data from wild-type cardioids, color-coded by cluster identity (left) or cardioid type (right). Data were downsampled to 900 cells per sample. Cluster identities are defined by expression of markers, with some examples shown on the bottom: *TNNT2* (CMs), *COL3A1* (fibroblasts), *ERG* (endothelial cells), *IRX4* (LV CMs), *IRX1* (RV CMs), *NR2F2* (atrial CMs), *KRT19* (epithelial cells), and *POU5F1* (pluripotency). (B) Cluster label transfer from the reference dataset GSE106118.

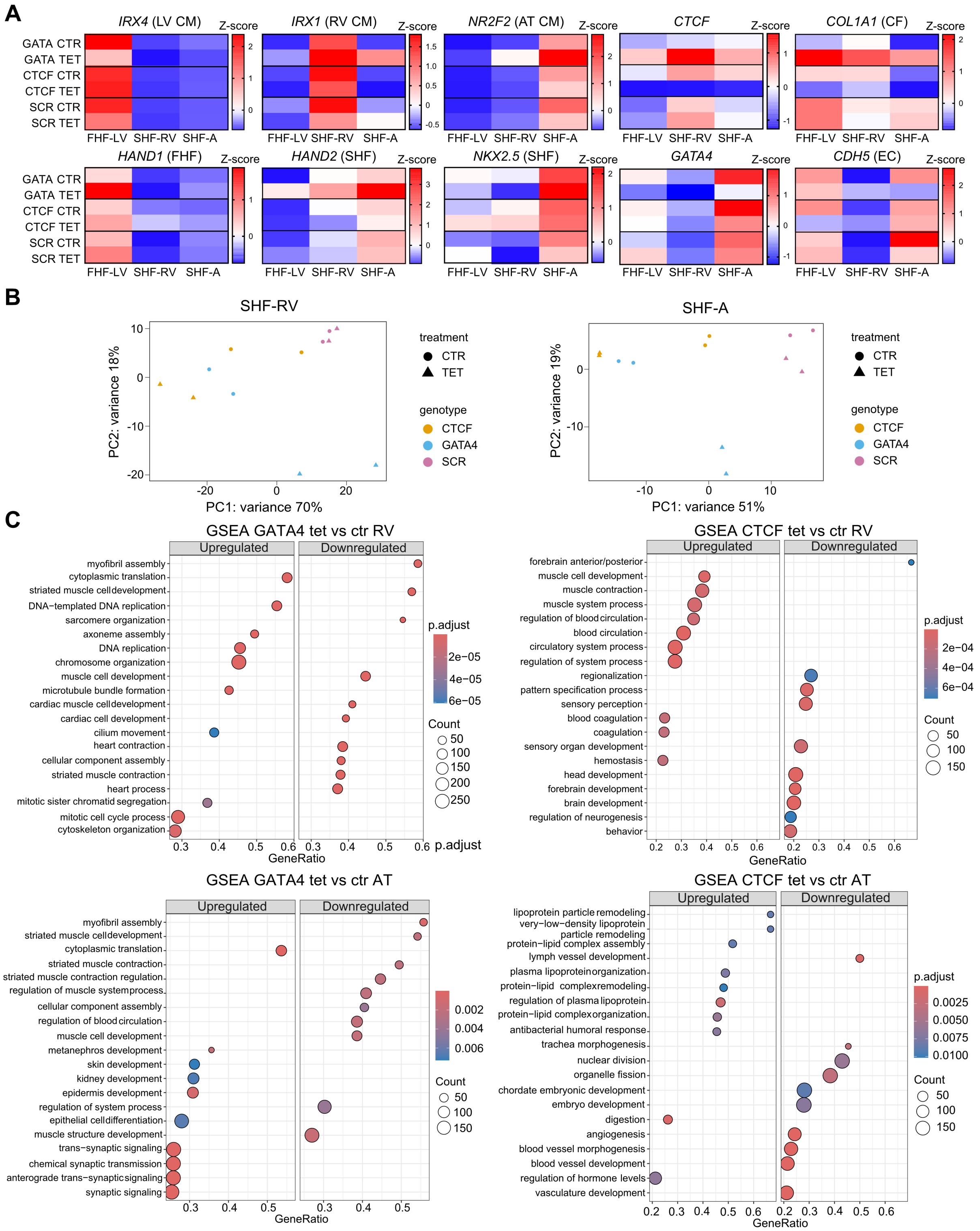


#### Figure S5. Inducible knockdown during SHF cardioid differentiation

Legend on the next page

#### Figure S5. Inducible knockdown during SHF cardioid differentiation (continued)

(A) RT-qPCR of chamber- and cell type–specific markers in day 7.5 cardioids after GATA4 or CTCF KD. n = 3 differentiations; average z-scores are shown.

(B) PCA of bulk RNA-seq data from SHF-RV and SHF-A cardioids after GATA4 or CTCF KD.

(C) GSEA on biological processes of GATA4 tet vs ctr and CTCF tet vs ctr in SHF-RV and SHF-AT day 7.5 cardioids. Data form bulk RNA-seq. N = 2 differentiations.

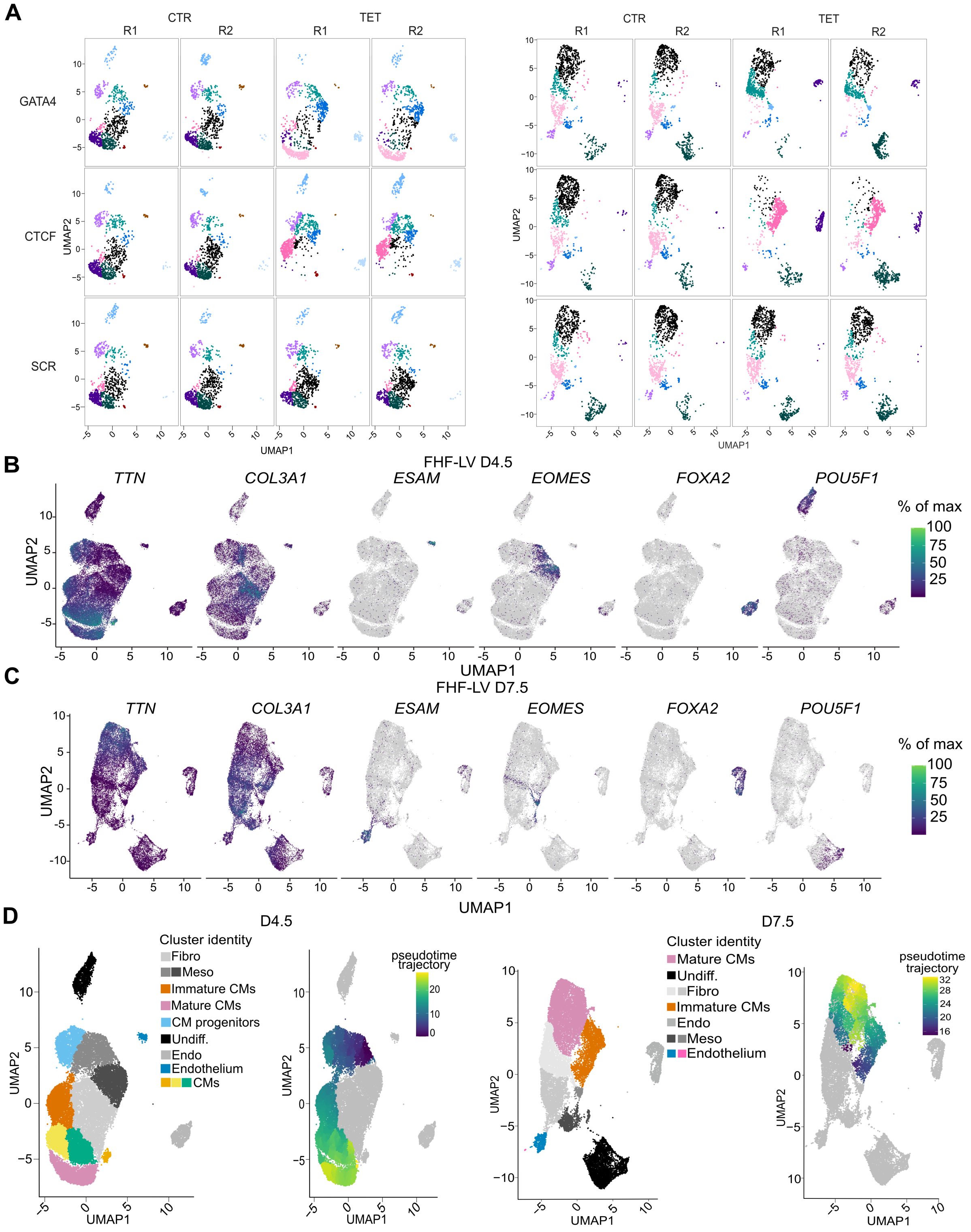


### Figure S6. Single-cell RNA-seq of differentiating inducible knockdown LV cardioids

Legend on the next page

#### Figure S6. Single-cell RNA-seq of differentiating inducible knockdown LV cardioids (continued)

(A) UMAP representation of scRNA-seq data from day 4.5 and 7.5 LV cardioids, color-coded by cluster identity. Biological replicates are shown side by side to illustrate reproducibility.
(B) Marker expression at day 4.5. *TTN*: CMs; *COL3A1*: fibroblasts; *ESAM*: endothelial cells; *EOMES*: mesoderm; *FOXA2*: endoderm; *POU5F1*: pluripotency.
(C) Marker expression at day 7.5.
(D) Cluster identity and pseudotime trajectories for CM clusters at day 4.5 and 7.5.

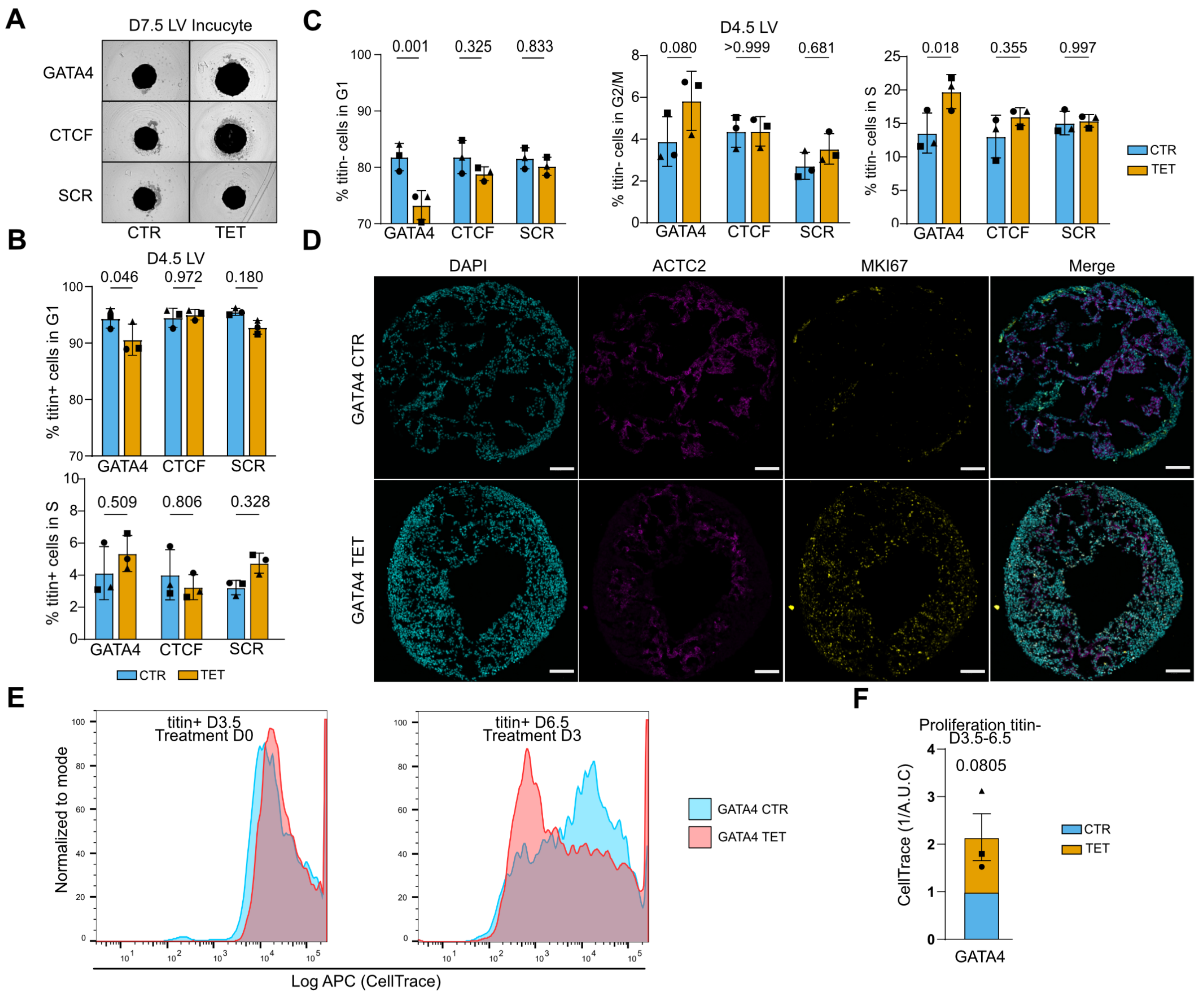


#### Figure S7. Proliferation dynamics in inducible knockdown LV cardioids

(A) Representative images of LV cardioids at day 7.5.

(B) Quantification of cell cycle distribution in titin$^+$ cells at day 4.5 LV cardioids, based on EdU incorporation and DNA content. n = 3 differentiations, p-values by two-way RM ANOVA with Holm–Sidak corrected pairwise multiple comparisons.

(C) As in panel B, but for titin$^-$ cells.

(D) Immunofluorescence of day 4.5 LV cardioids stained for $\alpha$-actinin 2 (ACTC2), nuclei (DAPI), and Ki-67 (MKI67). Scale bar: 100 µm.

(E) Representative flow cytometry plots of CellTrace signal in titin$^+$ cells at labeling (day 3.5) and three days later in GATA4 CTR and TET conditions.

(F) Proliferation assay with CellTrace dye. Cardioids were labelled at day 3.5 and analyzed by flow cytometry at day 6.5. Data show fold dilution of CellTrace signal in titin$^-$ cells, normalized to baseline (day 3.5). n = 3 differentiations, p-value by one-sample t-test against $\mu$ = 1.

#### Supplemental Tables

| Sequence | Top oligo (5'-3') | Bottom oligo (5'-3') |
|---|---|---|
| GATA4 shRNA | GATCCCGCCAGAGATTCTGCAAC ACGAACTCGAGTTCGTGTTGCAG AATCTCTGGTTTTTTG | TCGACAAAAAACCAGAGATTCTG CAACACGAACTCGAGTTCGTG TTGCAGAATCTCTGGCGG |
| CTCF shRNA | GATCCCGTATGATTTCCCATCGA CATTTCTCGAGAAATGTCGATGG GAAATCATATTTTTTG | TCGACAAAAAATATGATTTCCCAT CGACATTTCTCGAGAAATGTCG ATGGGAAATCATACGG |
| SCR shRNA | GATCCCGCGCGATAGCGCTAATA ATTTCTCGAGAAATTATTAGCG CTATCGCGCTTTTTTG | TCGACAAAAAAGCGCGATAGCG CTAATAATTTCTCGAGAAATTAT TAGCGCTATCGCGCGG |

**Table S1. Oligonucleotides for molecular cloning**

| PCR | Primer name | Primer sequence (5'-3') | Thermal protocol |
|---|---|---|---|
| CBS1 genotyping | CBS1_FW<br>CBS1_REV | AAACGTCGGTGCTTGAGTTG<br>TAGAAAGCACCGTATGCCGTC | Tann: 63; Ext time: 1'30" |
| CBS6 genotyping | CBS6_FW<br>CBS6_REV | AAACGTCGGTGCTTGAGTTG<br>TAGAAAGCACCGTATGCCGTC | Tann: 63; Ext time: 1'30" |
| AAVS1 WT locus | AAVS1_FW<br>AAVS1_REV | CTGTTTCCCCTTCCCAGGCAGGTCC<br>TGCAGGGGAACGGGGCTCAGTCTGA | Tann: 65; Ext time: 1'30" |
| AAVS1 5' junctional PCR | AAVS1_FW<br>PURO_REV | CTGTTTCCCCTTCCCAGGCAGGTCC<br>TCGTCGCGGGTGGCGAGGCGCACCG | Tann: 65; Ext time: 1'30'' |
| Random plasmid integration | PLASMID_FW<br>PLASMID_REV | ATGCTTCCGGCTCGTATGTT<br>TGAGGAAGAGTTCTTGCAGCTC | Tann: 60; Ext time: 1'30'' |

**Table S2. Oligonucleotides for genotyping**

| Gene name | Primer forward sequence (5'-3') | Primer reverse sequence (5'-3') |
|---|---|---|
| *GATA4* | AATCTAAGACACCAGCAGCTCCTTC | CATGGCCAGACATCGCACT |
| *CTCF* | ACGTCGGAATACCATGGCAAG | GGCTCTGGCTCAGGTTCAAT |
| *TBX5* | TCCAGAAACTCAAGCTCACC | TGGCAAAGGGATTATTCTCA |
| *NKX-5* | GAGCCGAAAAGAAAGCCTGAA | CACCGACACGTCTCACTCAG |
| *HAND2* | TCCAAAATCAAGACCCTGCG | GCTTTTCAAGATTTCGTTCAGC |
| *TTN* | GTAAAAAGAGCTGCCCCAGTGA | GCTAGGTGGCCCAGTGCTACT |
| *TNNT2* | TCAAAGACAGGATCGAGAGACG | CTGGATGTAACCCCCAAAATGC |
| *IRX4* | TCTACTGCCCGGTCTACGAG | GGAATCAAAGCTGTTCAGCGAG |
| *IRX1* | CGGTACGCGACAACTCTCTG | GACCCTTAATCAGGCGGACG |
| *NR2F2* | TCCTGTTCACCTCAGATGCC | AGCTTTCCGAATCTCGTCGG |
| *CHD5* | TTGGAACCAGATGCACATTGAT | TCTTGCGACTCACGCTTGAC |
| *COL1A1* | GGAGGAATTTCCGTGCCTGG | CAATCCTCGAGCACCCTGA |
| *PBGD* | GGAGCCATGTCTGGTAACGG | CCACGCGAATCACTCTCATCT |
| *HAND1* | TCAAAGACGCACTCTTCCAC | GTGCAGCGACAAAAAGAAAA |
| *RPLP0* | GGCGTCCTCGTGGAAGTGAC | GCCTTGCGCATCATGGTGTT |
| *NEBL* | TATGCTCTCTGAAAAGGCGAG | GATAGGCTCCGGGAAGAACAG |
| *ACTN2* | TTGACAGGAGGAAGAATGGCC | ATAATGCGGGCAAATTCGGC |
| *RYR2* | CTAATGTCTGGGTGGGCTGG | TGCTGCGTTTGATGCTTTCA |
| *CTNNA3* | CACCGTGGAGAAGCTACTGG | TCAGAGCTTCACTTTCTTTGCG |
| *SLC8A1* | AGACCTGGCTTCCCACTTTG | TGGCAAATGTGTCTGGCACT |
| *LMO7* | CAGTTGAAGCAGGTAGCCCA | CCATAGCGCCTCACATCCAT |
| *CCDC141* | GTTGCGCTTTCTACGACGAC | CCCGATCTTCCAAAGCCTTG |
| *MEF2A* | GCAATGCAGGTGGGATGTTG | GAGGGGGAGACTTTGTAGGC |
| *TGFB2* | CTTTGGATGCGGCCTATTGC | TAAGCTCAGGACCCTGCTGT |
| *CAMK2D* | CAACTATGCTGGCTACAAGGAA | CCCAAAGCTTCAGGTTCAAA |
| *CDH2* | AGAAGAAGACCAGGACTATGACTTG | TCATTGTCAGCCGCTTTAAGG |
| *ACTA2* | CCCGCCCAGAAACTAGACAC | GTCGCCCACGTAGGAATCTT |
| *RGS5* | CTCCTCCAGAAGCCAGACTC | AAGTCCATAGTTGTTCTGCAGG |
| *HPRT* | TGACACTGGCAAAACAATGCA | GGTCCTTTTCACCAGCAAGCT |

**Table S3. Oligonucleotides for RT-qPCR**

| Region of interest | Primer forward sequence (5'-3') | Primer reverse sequence (5'-3') |
|---|---|---|
| TTN CBS1 | AGCAAGTGTAAGGGGAAATGGA | CTGAACCTGGTTTTCCACCA |
| TTN CBS2 | AGCAATAATTCTGGGGATTGGC | TAGGGCAATACTGCCACCTAGA |
| TTN CBS3 | GCCTCCAAATGACACTGACTC | AAGGTCTCCAGTGCCACCTA |
| TTN CBS4 | AGGCAATTCCACCTCTTGACC | AATTTCACTCTGCTGACCCGA |
| TTN CBS5 | AGCGGTTATACCTCTAGGTGG | TGACTGAGGAGCCTGAAGAA |
| TTN CBS6 | TGTGGAGGGCGTAGATTGGA | CAGGAGTGGTTCCTCAGCATT |
| Neg ctrl 1 | GGTCCACCTGAAATGCCAAC | AGTGCTTAGGCACAGGCAAA |
| Neg ctrl 2 | ATGACCCTTCAAAACACACATC | CAGGCTAATTTTAAGAGGCCCAT |
| Neg ctrl 3 | CATGGATGGTCTTCTGAGGCA | TTCTTTCCCATAGTGTTGCCA |
| Neg ctrl 4 | CCTGCTGGTGAAGACACAAA | ACACCATTCTAAGCTCACAGGC |
| Pos ctrl 1 | GAAGTCCCAGCCTCAGGTTT | GCGAGACGCTTGTCCTACAG |
| Pos ctrl 2 | GAGGAGGCGCTGTAGGACAA | ACCGTCGACCCTTGTTCGAG |

**Table S4. Oligonucleotides for ChIP-qPCR**
